# The Homeodomain-interacting protein kinase Hipk promotes apoptosis by stabilizing the active form of Dronc

**DOI:** 10.1101/2025.06.12.659078

**Authors:** Juan Manuel García-Arias, Rafael Alejandro Juárez-Uribe, Luis Alberto Baena-López, Ginés Morata, Ernesto Sánchez-Herrero

## Abstract

Members of the evolutionarily conserved homeodomain-interacting protein kinase (Hipk) family play a critical role in regulating essential signalling pathways involved in growth, differentiation, and apoptosis. While vertebrates have multiple *hipk* genes, *Drosophila* contains a single *hipk* ortholog, what facilitates functional analysis. We find that *hipk* is necessary for the stabilization of the initiator caspase Dronc, thus enhancing the two Dronc activities in apoptotic scenarios: the induction of the caspase cascade, and the reinforcement of JNK signalling pathway. Conversely, our data suggest that Dronc also raises the expression levels of Hipk, thereby reinforcing the apoptotic response. These findings significantly enhance our understanding of caspase regulation and position Hipk as a promising target for modulating caspase activity in a variety of biological contexts.

## Introduction

Apoptosis, one of the most prevalent forms of programmed cell death, is a conserved phenomenon by which cells are eliminated through the function of an evolutionary conserved group of cysteine proteases, termed caspases, that dismantle the protein substrates and cause cell death (reviewed in Fuchs and Steller, 2011). Apoptosis can take place during normal development, like in the sculpting of Drosophila embryonic cephalic structures (Lohmann et al., 2002) or the elimination of the interdigital membranes in vertebrates (Jacobsen et al., 1996), or be triggered by stress or tissue damage (reviewed in Baonza et al. 2022).

Because of the simplicity of its genetic system and its sophisticated genetic technology, *Drosophila* is a useful model to analyse the regulation of apoptosis. Within *Drosophila,* the wing imaginal disc is especially convenient for the analysis of the apoptosis mechanisms, since little developmentally programmed apoptosis exists, but still shows a robust apoptosis induction in response to stressors like ionizing radiation (IR), heat shock, and others (Perez-Garijo et al., 2004; Verghese and Su, 2016). Moreover, apoptosis can experimentally be manipulated by driving the expression of numerous members of the apoptotic cascade including the pro-apoptotic genes (Grether et al., 1995; Chen et al., 1996; White et al., 1996; Smith-Bolton et al., 2009; Martin et al., 2017).

As in mammals, the apoptotic pathway in *Drosophila* engages pro-apoptotic genes, initiator and effector caspases and natural inhibitors of apoptosis such as Diap1 (Steller, 2008). An important feature of the apoptotic pathway in *Drosophila* is that it includes a feedback amplification loop (Supplementary Fig. 1), necessary for the full apoptotic response (Shlevkov and Morata, 2012). This loop involves the Jun N-Terminal Kinase (JNK) pathway, a versatile signalling pathway implicated in a number of biological processes (reviewed in Weston and Davis, 2007) including apoptosis in response to stress (McEwen and Peifer, 2005; McNamee and Brodsky, 2009). Upon irradiation there is an initial apoptotic stage, triggered by the DNA damage response pathway, which induces the function of the initiator caspase Dronc and the effector caspases Drice and Dcp1 (reviewed by Baonza et al., 2022). A second phase, consolidating the apoptotic response, appears to rely on a Dronc-dependent stimulation of the JNK signalling pathway (Wells et al., 2006; Shlevkov and Morata, 2012; Fogarty et al., 2016; Pinal et al., 2019). Despite intensive research, the molecular crosstalk between major signalling pathways, such as the JNK pathway, and the core apoptotic machinery, remains poorly understood.

A group of factors involved in the regulation of JNK signalling are members of the conserved Homeodomain-interacting protein kinase family, encoded by the *hipk* genes (reviewed in Schmitz et al., 2014; Blaquiere and Verheyen, 2017). While vertebrates possess four *hipk* members (*hipk1-4*), *Drosophila* only contains one, what facilitates the experimental analysis of Hipk function. The *Drosophila hipk* gene shows the highest homology with the vertebrate *hipk2* (Choi et al., 2005), which encodes a protein known to interact with many transcription factors and to regulate a variety of processes, including transcriptional regulation, chromatin remodelling, cell proliferation and apoptosis (Rinaldo et al., 2007). The *hipk* gene of *Drosophila* is also involved in the regulation of different processes by major pathways like Notch (Lee et al., 2009a), Wg (Lee et al., 2009b; Verheyen et al., 2012), Hippo (Chen and Verheyen, 2012; Steinmetz et al., 2021), JAK/STAT (Tettweiler et al., 2019), and JNK (Huang et al., 2011). Nevertheless, the molecular bases of these interactions are largely unknown, particularly in the context of apoptosis and JNK signalling regulation.

In this work we present evidence that the apoptotic activities of Dronc and the JNK signalling amplification are critically influenced by *hipk*. Specifically, our results indicate that to large extent these effects stem from the mutual ability of Hipk and Dronc to regulate each other’s activities in vivo.

## Results

In the following experiments we analyse the role of *hipk* in the implementation and regulation of developmentally programmed apoptosis.

### Developmentally programmed apoptosis requires *hipk* function

Previous data have indicated that *hipk* is required for the implementation of apoptosis in developmentally regulated scenarios in *Drosophila*. In particular, it has been shown that Hipk facilitates the removal of embryonic neurons and epithelial wing cells after adult hatching from the puparium (Link et al 2007). To further characterize the role of *hipk* during apoptosis, we have analysed the consequences of compromising *hipk* function in two developmental contexts showing intrinsic apoptosis: the fusion of the adult abdominal hemi-segments, and the rotation of the male genitalia.

During pupal development, polytene Larval Epidermal Cells (LECs) undergo cell death and are extruded from the epithelium. The elimination of LECs is tightly coupled with the proliferation of histoblasts that ultimately form the adult abdominal cuticle (Madhavan and Madhavan, 1980; Ninov et al., 2007; Bischoff and Cseresnyés, 2009; Nakajima et al., 2011). The dorsal histoblasts from the left and right sides meet at the midline to form a continuum epithelium in each abdominal segment (Fig. 1A). This process depends on apoptosis-mediated elimination of the LECs, as blocking apoptosis execution by overexpressing *p35* (Hay et al., 1994) under control of the LEC-specific *Eip71CD-Gal4* driver (Cherbas et al., 2003) results in lack or aberrant abdominal fusion at the midline (Kester and Nambu, 2011; Fig. 1B). Since JNK is an apoptosis inducer (McEwen and Peifer, 2005; McNamee and Brodsky, 2009; Pinal et al 2019), we also examined whether this pathway was necessary for the fusion of the adult hemi-segments. As shown in Figure 1C, inactivation of the JNK pathway in LECs compromised the fusion between abdominal hemi-segments. Interestingly, reducing *hipk* expression in these same cells with a validated *hipk*-RNAi construct (Chen and Verheyen, 2012) caused a weaker but comparable fusion defects to those observed blocking apoptosis or the JNK pathway (Fig. 1D).

**Figure 1.**
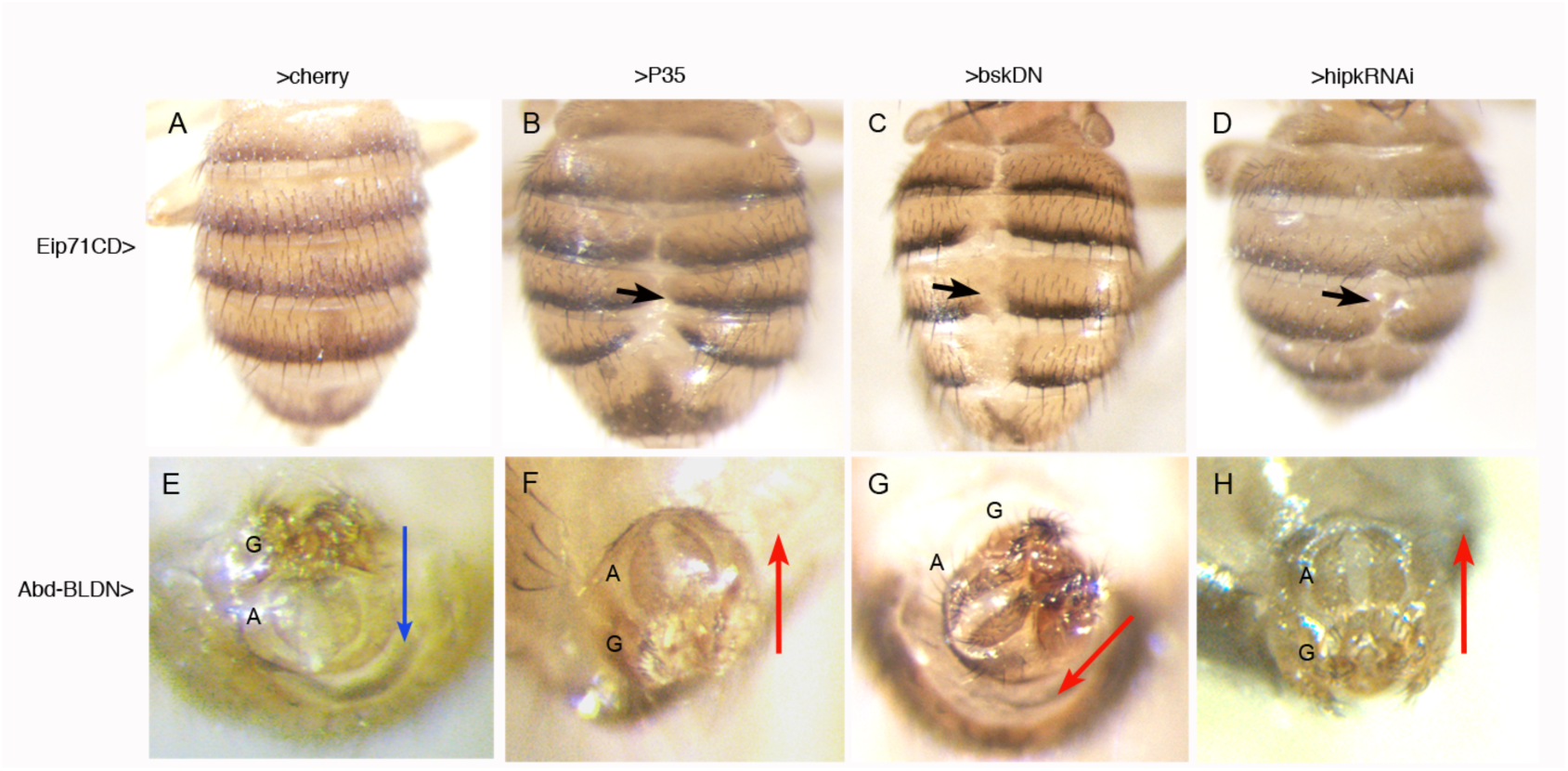
**The function of *hipk* is required for developmentally programmed apoptosis in the abdomen and terminalia** (A) Control *Eip71CD*-Gal4 UAS-*cherry* female dorsal abdomen showing continuous epithelium between left and right sides of each segment. (B) When the caspase inhibitor *p35* is expressed in the LECs (*Eip71CD-Gal4 UAS-p35* flies), some left and right dorsal tergites do not meet properly at the midline (arrow). (C) Suppression of JNK activity in the larval epidermal cells causes a similar phenotype. (D) When *hipk* expression is reduced in LECs (*Eip71CD-Gal4 UAS-hipk^RNAi^* flies) the phenotype is also similar, though weaker. 8-10 females were studied for each genotype, all showing a uniform phenotype. (E) Male genitalia and analia of a control *Abd-B^LDN^ UAS-cherry* fly, showing the wildtype disposition, genitalia (G) in the upper location and analia (A) in the lower one, as indicated by the arrow from genitalia to analia. (F, G) in *Abd-B^LDN^ UAS-p35* (F) o *Abd-B^LDN^ UAS-bsk^DN^* (G) males the normal disposition of the analia and genitalia is reversed or abnormal, the analia now being in an anterior or lateral location with respect to the genitalia, arrows. (H) In an *Abd-B^LDN^ UAS-hipk^RNAI^* male a similar abnormal phenotype is observed. 7-11 males were observed for each genotype, all with similar phenotypes. All the crosses were made at 25°C and the larvae transferred to 31°C to complete development.

Next, we examined the requirement of *hipk* function for the 360° rotation of the genital plate during early pupa (Gleichauf, 1936; Spéder et al., 2006; Hozumi et al., 2006). This process locates the genitalia in the upper and the analia in the lower position of the terminalia (Fig. 1E). This arrangement requires apoptosis in the eighth abdominal segment (A8) of the male genital disc before the genital plate starts to rotate, since there is impaired or absent rotation without apoptotic cell death (Abbott and Lengyel, 1991; Macías et al., 2004; Suzanne et al., 2010; Kuranaga et al., 2011). To manipulate the genetic expression within cells of the A8, we used an A8-specific Gal4 line (*Abd-B^LDN^*) to prevent cell death by forcing P35 activity or suppressing JNK in the A8 segment. In contrast to control flies only expressing the fluorescent protein Cherry (Fig. 1E), the overexpression of *p35* or a dominant negative form of *basket* (a key transducer of the JNK pathway) caused analia and genitalia rotation defects (Fig. 1F, G). Intriguingly, the expression of the *hipkRNAi* construct with the same driver also triggered similar phenotypes of abnormal disposition of genitalia and analia as those described upon suppressing cell death (Fig. 1H).

Taken together, these results establish that both the fusion of abdominal nests of histoblasts and the rotation of the male genitalia, require normal *hipk* function, likely by contributing to reach the apoptosis levels necessary to complete those processes.

### Experimentally induced apoptosis triggered by pro-apoptotic genes requires *hipk* activity

Next, we examined *hipk* role after induction of cell death by the pro-apoptotic gene *reaper* (*rpr*) (White et al., 1996; Martin et al., 2017). Rpr binds to the BIR domain of Diap1 facilitating its proteasomal degradation (Ryoo et al., 2002). This molecular interaction between Rpr and Diap-1 secondarily licenses for activation initiator and effector caspases, Dronc (Dorstyn et al., 1999) and Drice (Fraser and Evans, 1997), therefore inducing cell death (Goyal et al., 2000) (see Supplementary Fig. 1). To analyse the response to *rpr* induction of cells in which *hipk* function is compromised, we combined the transcriptional bipartite gene expression systems Gal4/UAS and LexA/LexO (Brand and Perrimon, 1993; Lai and Lee, 2006; see Material and Methods and drawings in Fig. 2A).

**Figure 2.**
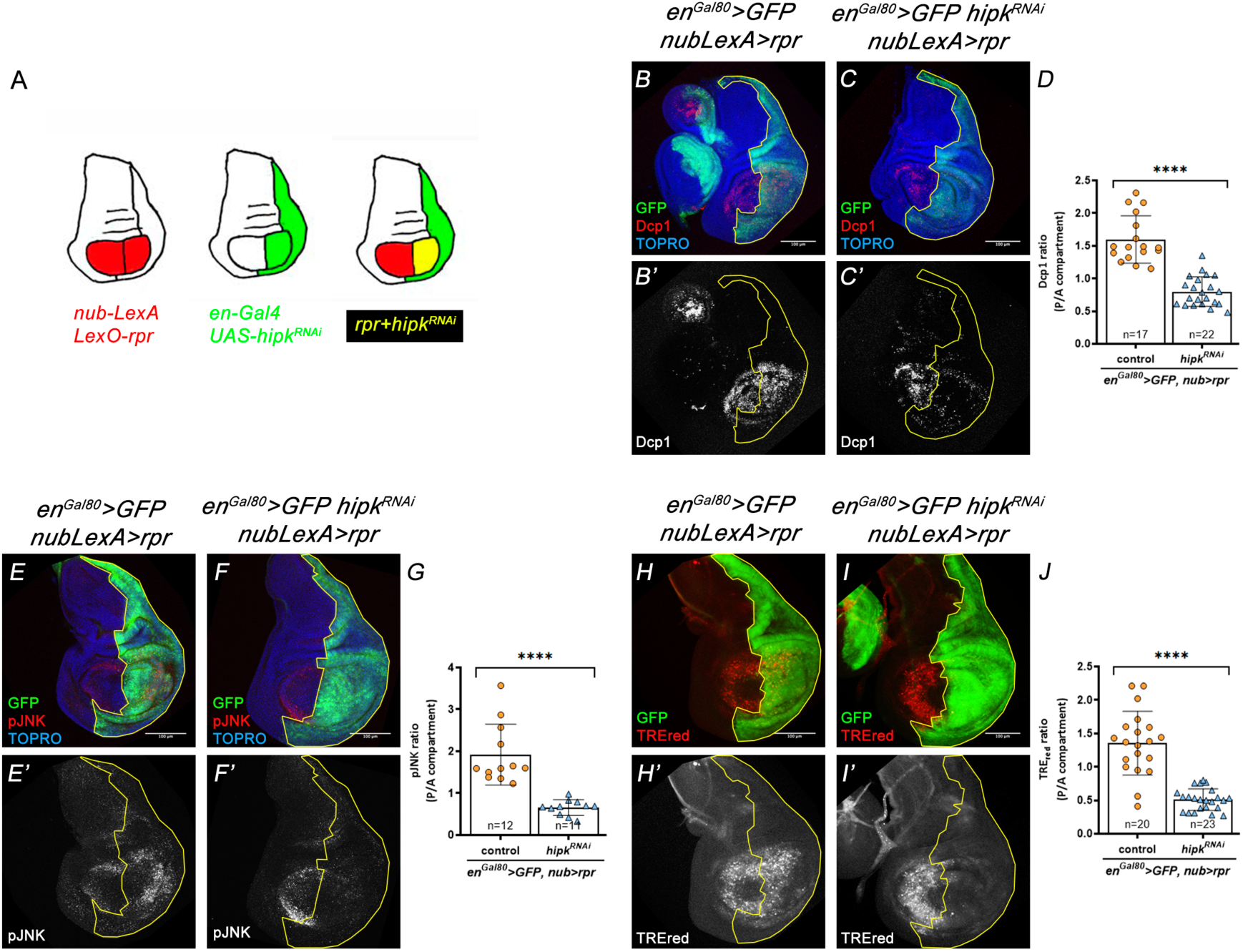
**The pro-apoptotic gene *rpr* requires the contribution of *hipk* for full apoptotic response and stimulus of JNK activity.** Genotypes on top of the panels. (A) Drawings illustrating the experiments. We have made use of the binary systems Gal4/UAS and LexA/LexO. The *en*-Gal4 driver directs GFP expression and reduces *hipk* levels only in the Posterior (P) compartment, thus discriminating between the Anterior (A) and the P compartments. The *nub*-LexA driver directs *rpr* expression (in red) in the wing pouch, which contains anterior and posterior regions. The combination of the two drivers permits to differentiate regions that contain only *rpr* expression (red), only *hipk* expression (green) or both (yellow). (B, B’, D) In *en^Gal80^ >GFP*; *nub-LexA>rpr* discs the entire P compartment is labelled with GFP, and *rpr* is expressed in the Nubbin domain. Staining with the marker Dcp1 (red) shows high apoptotic levels in the entire Nubbin domain, A and P compartments. (C, C’, D) In contrast, in *en^Gal80^ >GFP hipk^RNAi^ nub-LexA>rpr* discs there is a marked reduction of Dcp1 in the posterior Nubbin region. The images in E-F’, G and H-I’, J illustrate similar experiments demonstrating the effect of the loss of *hipk* function on JNK activity, monitored by the expression of the phosphorylated form of Jun (E-G) or by the TRE-red marker (H-J).

In control wing discs, forced expression of *rpr* in nubbin-expressing cells (in the wing pouch) induced strong cleaved Dcp-1 immunoreactivity and JNK activation—both established markers of apoptosis (Fig. 2B, B’, E-E’, H-H’). However, a reduction of *hipk* expression yielded a potent rescue of these features (Fig. 2C-C’, D, F-F’, G, I-I’, J). These results suggested that *hipk* is key for both *rpr*-induced JNK signalling and apoptosis.

The requirement of *hipk* function in the maintenance of JNK activity was further investigated in an experiment in which apoptosis was induced by IR but the execution of the apoptosis programmed was prevented overexpressing the effector caspase inhibitor *p35*. In this experimental setting, previous work (reviewed in Pinal et al., 2019) demonstrated that *dronc*-dependent activation of JNK and persistent proliferative signalling emanating from these cells induces wing imaginal discs overgrowth after irradiation. In line with these observations, irradiated discs expressing *p35* in wing disc posterior cells (P compartment) show a significant increase in size and ectopic activity of JNK, as indicated by the Mmp1 marker (Uhlirova and Bohmann, 2006), compared to non-irradiated control discs (Supplementary Fig. 2A, A’, B, B’, D, E). However, there was no overgrowth and limited JNK activation upon irradiation of P35 cells without *hipk* expression (Supplementary Fig. 2C, C’, D, E).

### *hipk* does not primarily exert its effect through *Diap1*

In addition to its effect on apoptosis and JNK activity, we found that the suppression of *hipk* function in *rp*r-expressing cells caused accumulation of the Diap1 protein (Fig. 3A, A’, B, B’, C), suggesting a role of *hipk* in the *rpr*-mediated degradation of Diap1 (Goyal et al., 2000). This result also suggested that the diminution of the apoptosis levels observed in the absence of *hipk* function could be due to maintenance of high levels of Diap1; the Diap1 protein has a key role in preventing the cleavage and subsequent activation of caspases (Meier et al., 2000), thereby the lack of Diap1 results in massive apoptosis (Hay et al., 1995; Wang et al., 1999; Goyal et al., 2000; Lisi et al., 2000; Ryoo et al., 2002). To test this, we compromised *diap1* expression in the P compartment by using an effective RNAi construct (Ryoo et al., 2004). In control discs, in which we reduced *diap1* levels, we detected consistent elevation of apoptotic marker (Fig. 3D, D’, F). In contrast, a significant downregulation of apoptosis markers was observed by concomitantly reducing *diap1* and *hipk* (Fig. 3E, E’, F). These important result rule out the hypothesis that *hipk* downregulates apoptosis by increasing Diap1 levels, and indicate that Hipk acts downstream of *diap1*, possibly facilitating the activation of the caspase pathway.

**Figure 3.**
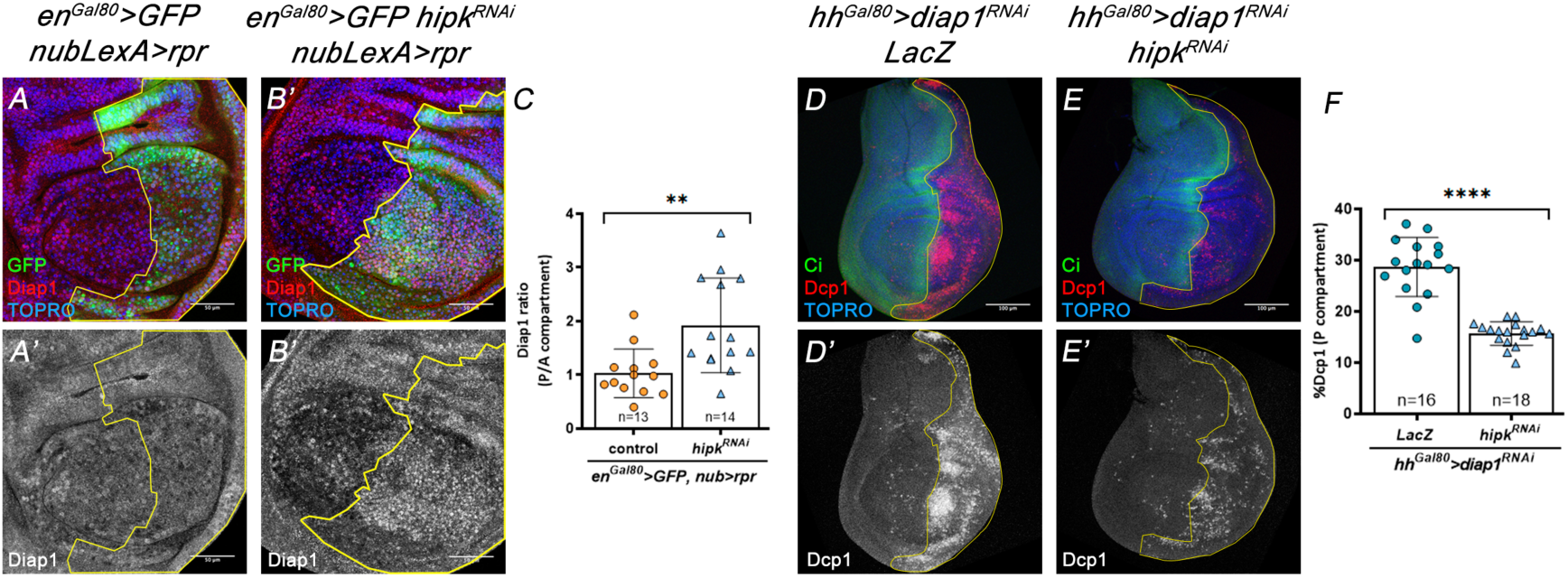
**Functional interactions between *hipk* and *diap1*.** Genotypes on top of the panels. (A-B’, C) Reduction of *hipk* function in the posterior region of the Nubbin domain of discs of genotype *en^Gal80^ >GFP hipk^RNA^ nub-LexA>rpr* (B, B’) causes an increase in the amount of Diap1 protein in the P compartment, something not observed in control discs (*en^Gal80^ >GFP nub-LexA>rpr*) (A, A’), as indicated by the levels of anti-Diap1 antibody. Quantification in C. (D, D’) Suppression of *diap1* in the P compartment of *hh^Gal80^* >*diap1^RNAi^ lacZ* discs causes a strong apoptotic response, as indicated by the accumulation of the Dcp1 caspase (red). The A/P boundary is delineated by the expression of Ci, an A compartment marker. (E, E’) Compromising *hipk* function by RNA interference in the P compartment of *hh^Gal80^* >*diap1^RNAi^ hipk^RNAi^* larvae results in a significant decrease of Dcp1 levels. Quantification in F.

### *hipk* promotes apoptosis mainly by stabilizing the active form of Dronc

To explore the hypothesis of a functional interaction between Hipk and the caspase cascade, we have analysed the effect of the lack of *hipk* on the apoptosis induced by the direct activation of caspases. We first checked that, as expected, the lack of *hipk* function does not affect the normal very low levels of apoptosis in the wing disc (Fig. 4A-B’, I). We then conducted three experiments that overexpressed *dronc*, *drice*, or both together. To overexpress *dronc* and *drice* together we capitalized on a UAS construct in which *dronc* and *drice* cDNAs were concomitantly overexpressed (see Methods). This combined overexpression induced prominent apoptosis in the P compartment of wing discs (Fig. 4C, C, I’). However, such apoptotic response was drastically rescued by downregulating *hipk* (Fig. 4D, D’ I). The single overexpression of either *dronc* or *drice* also induced apoptosis, though to a lesser scale (Fig. 4E, E’, G, G’, J, K). Interestingly, in these experiments Dronc-induced apoptosis, but not Drice-induced apoptosis, was rescued by limiting *hipk* expression (Fig. 4F, F’, J, H, H’, K). Altogether, these data indicated that *hipk* sustains the apoptotic response by likely acting at the level of the initiator caspase Dronc.

**Figure 4.**
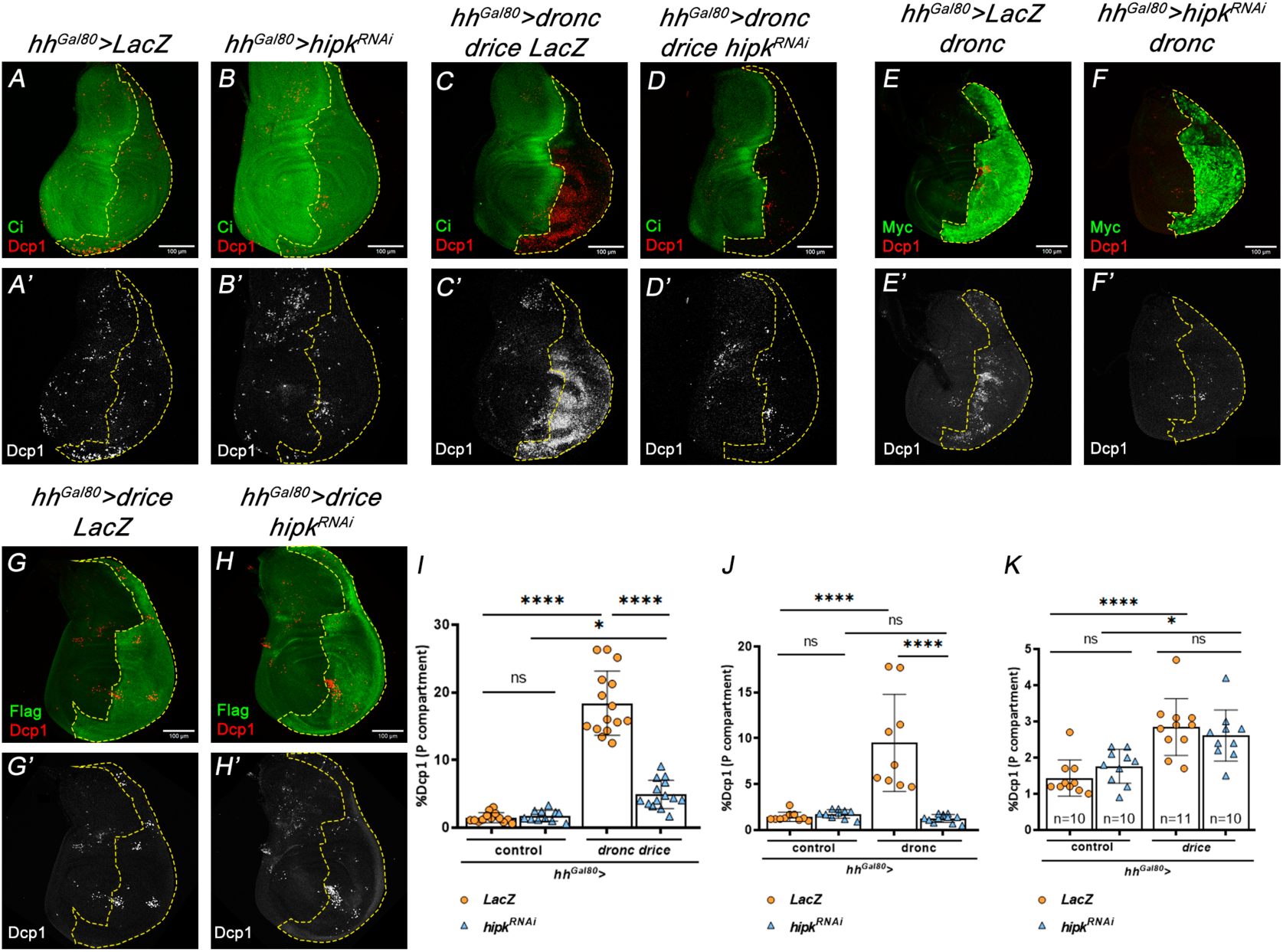
**Effect of *hipk* down-regulation on apoptotic levels induced by Dronc and Drice overexpression.** Genotypes on top of the panels. (A-B’) The amount of apoptosis (Dcp1, red) in control *hh>lacZ* (A, A’) and *hh>hipk^RNAi^* (B, B’) discs is similarly low. The Ci antibody marks the anterior compartment. Quantification in I. (C-D’) The joint overexpression of *dronc* and *drice* in the P compartment results in high levels of apoptosis (C, C’), as indicated by Dcp1 staining, but these are drastically reduced by compromising *hipk* function (D, D’). The Ci antibody marks the anterior compartment. Quantification in I. (E-F’) Overexpression of *dronc* in the P compartment (marked my antibody against the Myc tag) shows a moderate increase of apoptosis (E, E’), which is suppressed by compromising *hipk* function (F, F’). Quantification in J. (G-H’) Overexpression of *drice* (marked by an antibody against the Flag tag) causes a slight increase of apoptosis (G, G’), which is not affected by reducing *hipk* activity (H, H’). Quantification in K.

Our previous experiments took advantage of newly generated transgenic lines expressing tagged versions of Dronc and Drice (see Methods) suitable to assess their protein stability upon modulating Hipk expression levels. Specifically, Dronc is fused in-frame with a Myc tag and a modified GFP that only fluoresces upon Dronc-mediated cleavage (see Methods and relevant references). This dual-tagging system enables simultaneous detection of both total Dronc protein via anti-Myc immunolabelling and its activation via GFP cleavage. Drice, in turn, was tagged at the C-terminus with a Flag epitope. Remarkably, posterior cells overexpressing Dronc and Drice displayed strong Myc immunolabelling and GFP signal, indicating robust Dronc activation (Fig. 5A, A’’, C, D). However, a dramatic reduction of both Dronc protein levels and activation was detected in Hipk-deficient cells (Fig. 5B, B’’, C, D). A similar, albeit milder, reduction was observed in cells overexpressing Dronc with limited Hipk levels (Fig. 5E-F’’, G, H). In contrast, the detection of Drice through Flag immunostaining remained unaltered in cells with reduced Hipk (Fig. 5I-J’, K).

**Figure 5.**
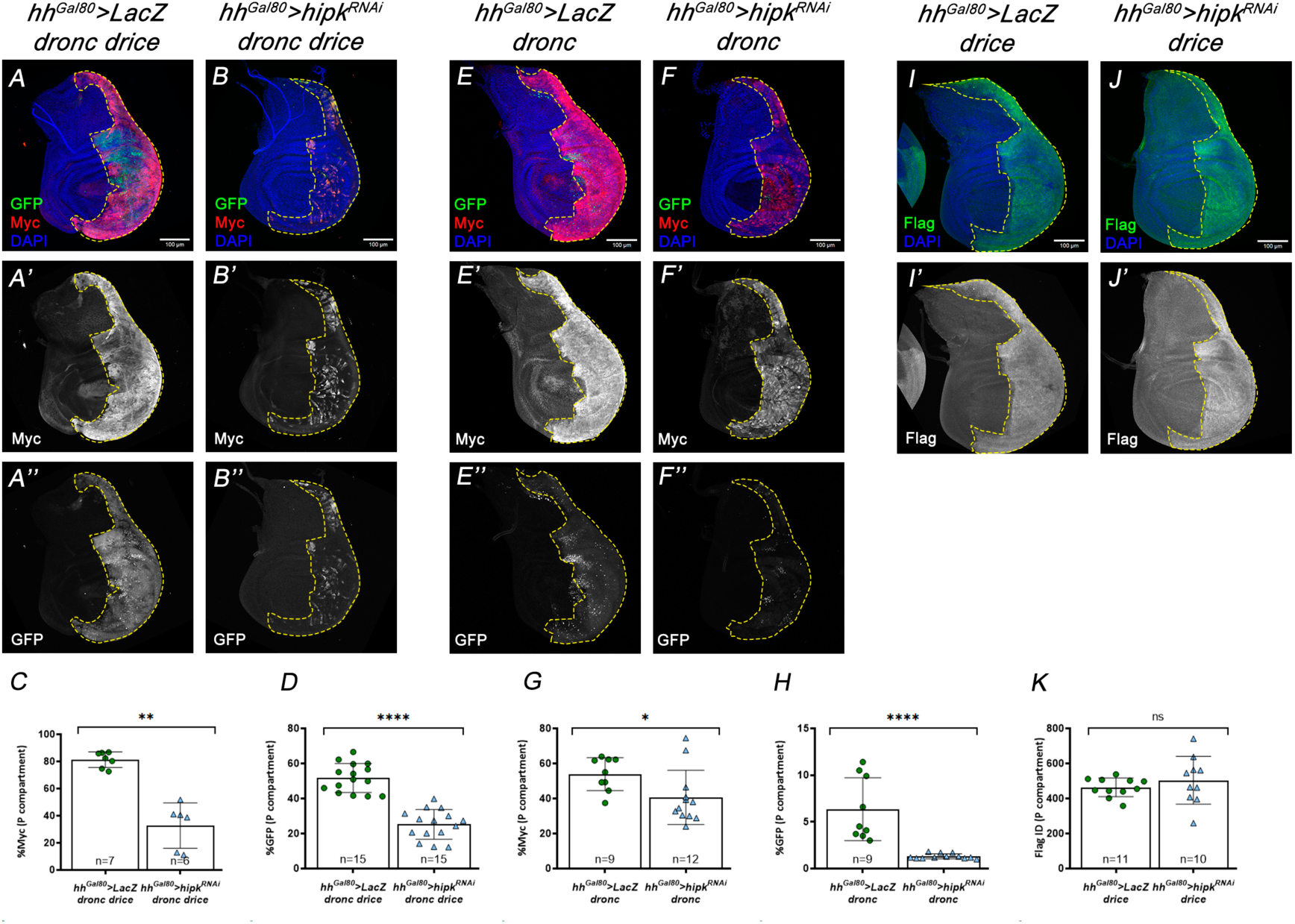
**Role of Hipk in the stabilization of the Dronc protein.** Genotypes on top of the panels. (A-B’’) Joint overexpression of *dronc* and *drice* cause an accumulation of total Dronc protein (A, A’) in the P compartment, as indicated by anti-Myc, as well as of active Dronc protein (A, A’’), as shown by GFP staining, but the lack of *hipk* function substantially reduces both (B-B”). Quantifications in C, D. (E-F’’) After overexpression of *dronc* in the P compartment, there is also an accumulation of both types of proteins (E-E’’), whose levels are reduced (more clearly for the active one) when *hipk* activity is reduced (F-F”). Quantifications in G, H. (I-J’’) Overexpression of *drice* in the P compartment results in high levels of Drice protein (labelled with Flag) (I, I’), which are not altered by compromising *hipk* activity (J, J’). Quantification in K.

Together, these results strongly support the notion that Hipk preferentially and intrinsically enhances the stability of Dronc in its active conformation. Reinforcing this conclusion, lack of *hipk* activity effectively rescues the tissue overgrowth caused by *p35*-expressing cells upon irradiation, which, despite lacking effector caspase activity, still activate Dronc (Supplementary Fig. 2B-D).

Further support for the interaction between *hipk* and *Dronc* comes from experiments of *hipk* overexpression. As previously reported (Chen and Verheyen, 2012; Poon et al., 2012; Blaquiere et al., 2018), the excess of Hipk caused mild tissue overgrowths (Fig. 6A-B’, E) that were enhanced by the concomitant overexpression of the pro-apoptotic gene *hid* (Fig. 6C, C’, E). However, this phenotype was critically linked to Dronc activity, as it was rescued in a mutant background null for *dronc* (Fig. 6D, D’, E). Similarly, forced expression of *hipk* is sufficient to activate Dronc and cleaved Dcp-1 immunoreactivity, but the absence of *dronc* drastically reduces Dcp1 levels (Supplementary Fig. 3A-E).

**Figure 6.**
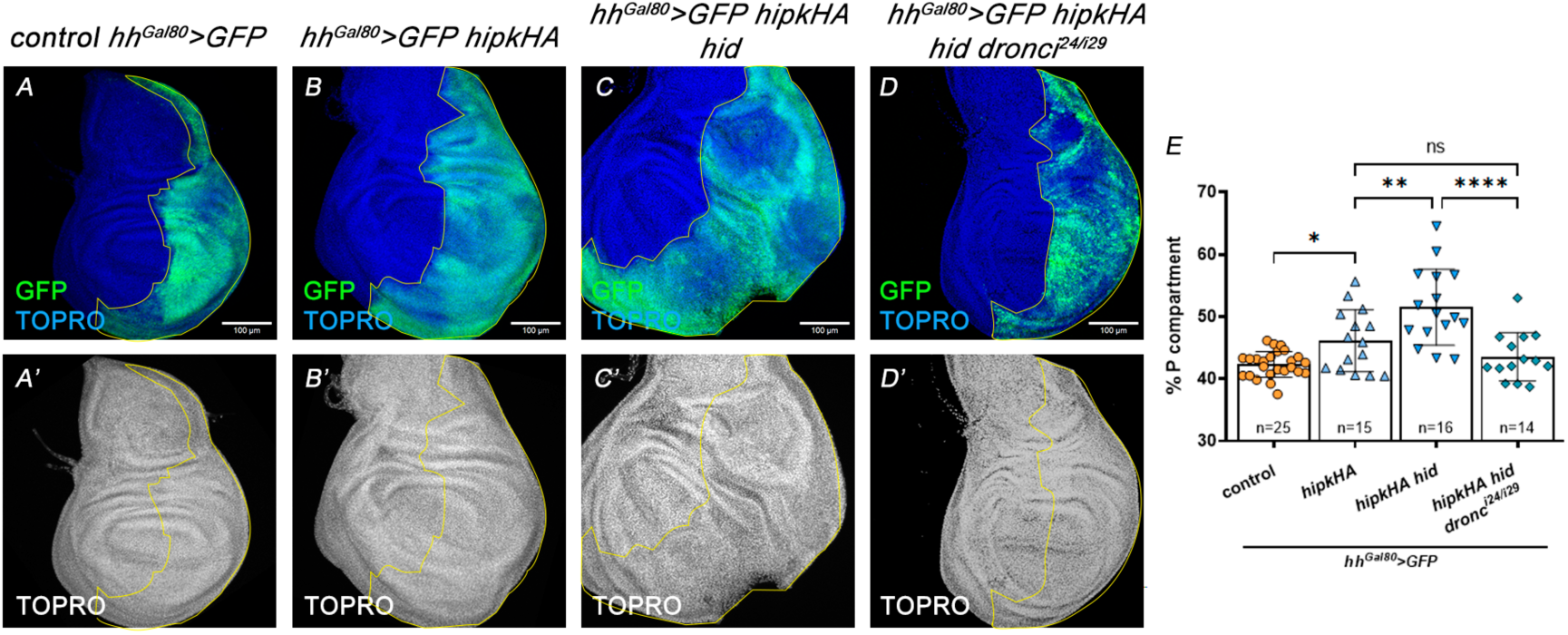
**The overgrowth caused by *hipk* is dependent on *dronc* function** Genotypes on top of the panels. (A, A’) Control *hh^Gal80^>GFP* wing disc in which the P compartment is labelled with GPP. (B, B’) Overexpression of *hipk*, using a UAS-*hipk-HA* construct expressed in the P compartment, causes a modest overgrowth of the compartment. (C, C’) The concomitant expression of *hid* and *hipk-HA* produces a larger increase in size of the compartment. (D, D’) When both *hid* and *hipk-HA* are expressed but in a *dronc* mutant background (*dronc^i24^*/*dronc^i29^*), the P compartment size is drastically reduced. Quantifications in E.

### Active Dronc also promotes Hipk protein stability

The observations above strongly suggest a determinant role of *hipk* to ensure the correct levels of apoptosis in either developmentally regulated or induced apoptosis. More specifically, our previous observations suggested that the Hipk protein preferentially affects the stability of Dronc in its active form. Since previous studies have shown that caspases can enhance Hipk2 activity in mammals (Gresko et al., 2006), we sought to investigate whether *Hipk* might, in turn, be regulated by Dronc. To his end, we first evaluated the Hipk levels in cells expressing *p35*, which cannot complete apoptosis but still activate Dronc after irradiation. Interestingly, in this experimental setting we found groups of *p35*-expressing cells showing significantly elevated levels of Hipk. A closer examination revealed that these cells also activated JNK signalling, as indicated by the Mmp1 upregulation (Fig. 7A-C’, E). This result suggested that ionising radiation could raise the amount of Hipk in Dronc-activating cells that fail to die. Notably, such upregulation of Hipk did not occur in irradiated discs in which the expression of pro-apoptotic genes was potently targeted by overexpressing a micro RNA against the proapoptotic genes Rpr, Hid and Grim (mirRHG) (Siegriest et al., 2010) (Fig. 7D, D’, F), despite the fact that the JNK pathway was still upregulated by apoptosis-independent JNK activation (Pinal et al., 2018) (Fig. 7D, D’, F).

**Figure 7.**
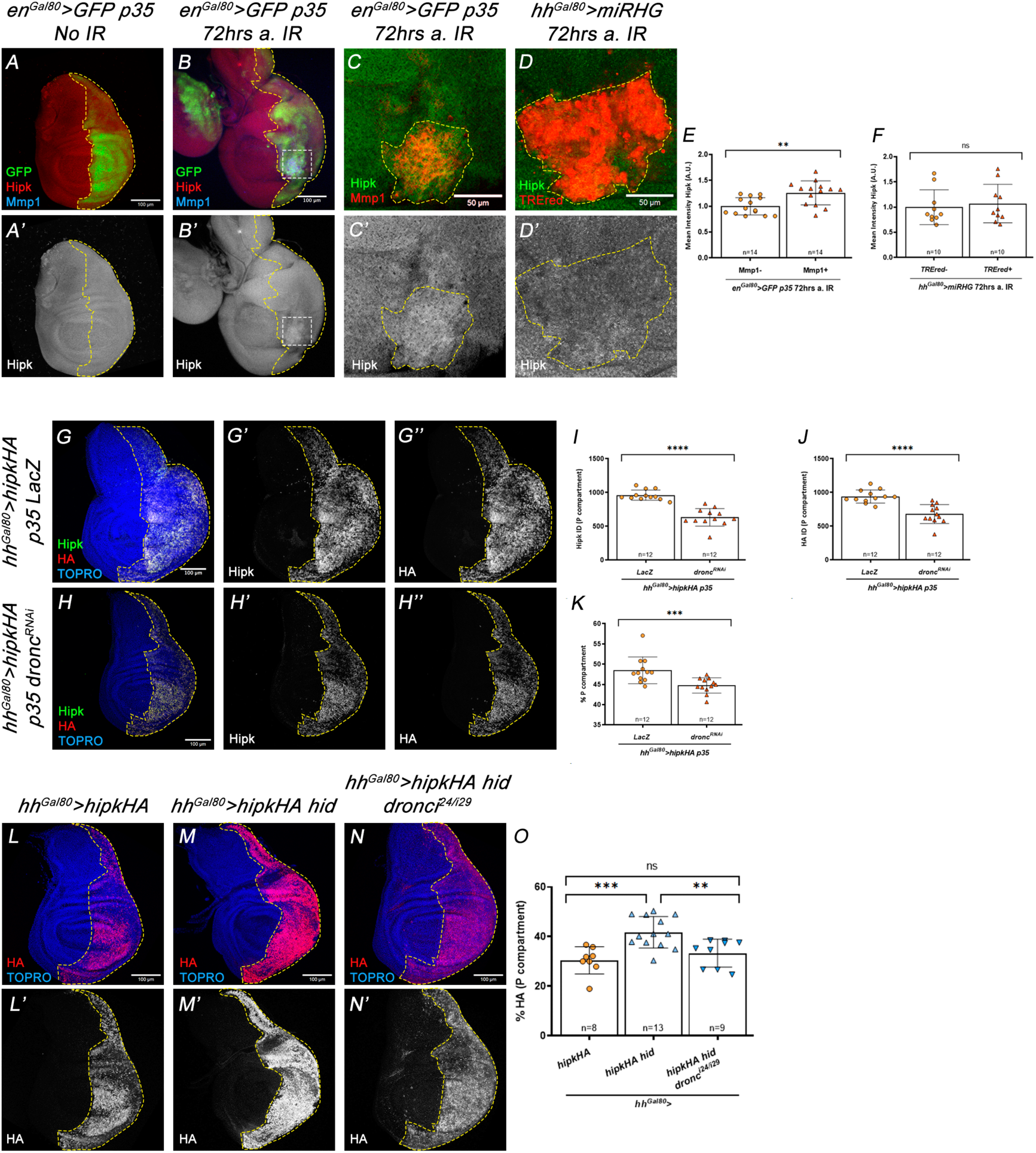
**Regulation of Hipk function and levels by pro-apoptotic genes and by *dronc*** Genotypes on top of the panels. (A, A’) In non-irradiated discs Hipk antibody expression is uniform in the wing disc (red); GFP expression (green) labels the posterior (P) compartment. (B, B’) After IR of *p35*-expressing wing discs, patches of higher Hipk expression are observed (inset). (C, C’) Magnification of the inset shown in B. The delineated area shows JNK activity, as indicated by expression of the Mmp1 marker (red), and increased levels of anti-Hipk signal. (D, D’) Portion of the P compartment of an irradiated disc in which activity of pro-apoptotic genes is suppressed by the presence of the *mirRHG* construct (Siegrist et al., 2010). The delineated patch shows JNK activity (TRE-red signal) but Hipk levels are not increased. Quantifications in E, F. (G-G’’) The overexpression of the *hipk* gene (*hipk-HA* construct) in the P compartment gives rise to an accumulation of the Hipk protein, as measured with the anti-Hipk and anti-HA antibodies. (H-H’’) If *dronc* expression is reduced in this genetic background, the amount of both anti-Hipk and anti-HA signals, as well as the size of the compartment, are clearly reduced. Quantifications in I, J, K. (L, L’) Forced expression of *hipk-HA* in the P compartment, showing anti-HA signal. (M, M’) The joint overexpression of *hipk-HA* and *hid* strongly increases anti-HA levels in this compartment, but the amount of this signal is drastically reduced in a *dronc* mutant background (N, N’). Quantifications in O.

These results argue for a role of the apoptotic program in elevating Hipk levels, but do not discriminate if this is a transcriptional or post-transcriptional effect, and do not single out Dronc as the key protein in this regulation. To solve these issues, we forced expression of a *hipk-HA* construct and quantified total Hipk and HA levels in *p35*-expressing cells with either normal or reduced Dronc expression. Intriguingly, absolute levels of Hipk-HA, detected using both anti-HA and a Hipk-specific antiserum were significantly reduced in Dronc-deficient cells with respect to controls (Fig. 7G-J). Consistently, these findings were correlated with a limited ability of Hipk overexpression to induce tissue overgrowth in cells without Dronc (Fig. 6C-E; 7K). To address whether Hipk upregulation was a consequence of impeding the completion of apoptosis via P35 or an effect connected to Dronc, we expressed the UAS-*hipk-HA* construct (along with *hid* but without *p35*) in either wild type or Dronc-deficient cells. The experiment revealed a significant upregulation of HA levels upon *hid* and *hipk-HA* co-expression and a strong reduction of HA levels when *dronc* was absent (Fig, 7L-O). Collectively, these findings support a reciprocal regulatory relationship: Hipk enhances the stability of Dronc, while active Dronc promotes the accumulation of Hipk.

## Discussion

In this report we provide compelling evidence that the Hipk plays a key role during the execution of apoptosis by stabilizing the active form of Dronc. Given the limited understanding of caspase regulation upon activation, our findings open a new avenue of research with significant implications for caspase biology. Conversely, we also show that Dronc promotes an increase in Hipk expression levels, further amplifying the apoptotic cascade and JNK activation. This intriguing observation likely reflects a positive feedback loop previously described between Dronc and the JNK pathway (Shlevkov and Morata, 2012).

### Hipk is a pro-apoptotic factor that stabilizes active Dronc

Hipk proteins are evolutionarily conserved serine/threonine kinases traditionally associated with fine-tuning transcriptional responses that affect various biological functions, such as cell proliferation, cell fate decisions, and apoptosis (Rinaldo et al., 2007; Blaquiere and Verheyen, 2017). Our results strongly suggest that *Drosophila* Hipk acts as a proapoptotic factor, as its reduced expression significantly diminishes apoptosis in either developmentally regulated or experimentally induced apoptotic contexts. The implication of *hipk* in developmentally regulated apoptosis has been reported previously (Link et al 2007) and we have confirmed this requirement in the left-right fusion of the abdominal hemisegments and the rotation of the male genitalia. In addition, we show that the activity of JNK is also necessary in both processes, thus pointing to a relevant role of JNK in developmental regulated apoptosis.

We have also demonstrated the involvement of *hipk* in the response to various pro-apoptotic stimuli. Our epistasis experiments show that the loss of Hipk function robustly suppresses apoptosis triggered by either overexpression of pro-apoptotic factors, the loss of cell death inhibitors such as Diap-1, the combined overexpression of initiator and effector caspases, or by initiator caspase Dronc alone. In contrast, cell death driven solely by effector caspase overexpression (e.g., Drice) remains largely unaffected by Hipk deficiency. All these experiments position Hipk activity at the level of the initiator caspase Dronc. In parallel, we found in our experiments that Hipk deficiency also compromises JNK activation in apoptosis-induced scenarios, thereby suggesting a Hipk-mediated control of the two Dronc activities: the induction of apoptosis through Dcp1 and Drice, and the amplification of apoptosis and JNK activity through the apoptotic loop (Shlevkov and Morata 2012).

In mammalian systems, Hipk2 also mediates apoptosis, but by direct phosphorylation of P53 (Hofmann et al., 2003; D’Orazi et al., 2002; Di Stefano et al., 2004; Dautz et al., 2007) and/or by facilitating the degradation of its inhibitor, MDM2 (Wang et al., 2001). In parallel, members of the Hipk family have been shown to potentiate JNK pathway activation (Hofmann et al., 2003) and apoptosis (Zhang et al., 2003; Sombroek and Hofmann, 2009) by antagonizing transcriptional repressors of the CtBP family. Thus, in Drosophila and mammals Hipk members regulate apoptosis and JNK signalling, although the molecules involved in such regulation may be distinct.

Our experiments indicate that Hipk plays a critical role in stabilizing Dronc, most notably, the active form of Dronc. Whereas Hipk moderately alter Dronc protein levels under basal, non-apoptotic conditions, in cells exposed to apoptotic stimuli Hipk is substantially required to sustain Dronc stability. The finding that loss of *hipk* causes a diminution of Dronc product may suggest that a primary cause of *hipk* phenotypes is precisely a reduction in the amount of active Dronc protein available to fulfil those roles, what results in partial suppression of Dronc function. Importantly, this regulatory mechanism may differ from those previously described. Thus, protein– protein interactions with Dark (Rodríguez et al., 1999), and Tango7 (D’Brot et al., 2013) have been shown to promote the assembly of protein complexes that enable efficient Dronc activation, while interaction with MyoID localizes Dronc to specific subcellular compartments (Amcheslavsky et al., 2018). Moreover, Hipk probably does not exert its pro-apoptotic function through its canonical role in modulating transcriptional regulation. Furthermore, our data raise the possibility that Hipk modulates the stability of active Dronc through phosphorylation—either directly or by influencing upstream regulators involved in its turnover. Such post-translational regulation would not be unexpected, as phosphorylation-based control of caspases has been reported in mammals (Brady et al., 2005; Martin et al., 2005; Allan and Clarke, 2009; Ge et al., 2024), and Dronc in *Drosophila* (Yang et al., 2010). Regardless of the ultimate molecular mechanism by which Hipk acts on Dronc, our findings clearly establish that Hipk enhances Dronc function and promotes apoptosis, reveal a novel regulatory pathway that modulates Dronc function, and broaden our current understanding of caspase biology.

### Hipk, JNK pathway and Dronc key players forming a positive apoptotic feedback loop

The Hipk protein interacts with different transcription factors and other molecules implicated in distinct biological operations (Blaquiere and Verheyen, 2017). The levels of *Drosophila* Hipk must be tightly regulated since both overexpression or loss of function of *hipk* can induce apoptosis (Chen and Verheyen, 2012). We have found that pro-apoptotic stimuli like IR cause an elevation of the amount of the Hipk protein, and this increment requires normal function of the apoptotic cascade. This process would ensure that there is a surplus of active Hipk necessary for activation of Dronc. More specifically, our results show Dronc is needed to maintain Hipk levels, which suggests a mutual interaction between Dronc and Hipk to reciprocally sustain their stability.

Interestingly, in mammalian cells, it has been reported that stress-induced activation of Caspase-6 leads to the proteolytic processing of Hipk2 (Gresko et al., 2006; Sombroek and Hofmann, 2009). Notably, this cleavage event removes an inhibitory C-terminal domain, generating a hyperactive kinase that further amplifies apoptosis. These and our own findings suggest that caspases could be evolutionarily conserved regulators of Hipk, capable of modulating either its protein abundance or activity. This mutual regulation between caspases and Hipk may be critical for amplifying the apoptotic response in diverse cellular contexts across evolution and could represent a targetable axis for future therapeutic interventions.

This mutual Dronc-Hipk interaction also impinges on activation of JNK signalling, a central, evolutionarily conserved regulator of apoptosis (Liu and Lin, 2005). In *Drosophila*, Hipk acts as a positive regulator of the JNK pathway in wing imaginal discs. Its activity is tightly regulated by SUMOylation, and upon loss of SUMO modification (e.g., through Smt3 knockdown), Hipk accumulates in the cytoplasm, enhancing JNK pathway activation and apoptosis (Huang et al., 2011). In vertebrates, Hipk proteins appear to function as key positive regulators of JNK signalling and c-Jun phosphorylation, through both direct and indirect mechanisms that are highly context-dependent (Hofmann et al., 2003; Zhang et al., 2003). Given Hipk’s known role in modulating diverse cellular processes, it is also tempting to speculate that, in addition to JNK signalling, Hipk may also regulate other non-apoptotic functions of Dronc, but further work is needed to validate this hypothesis.

In summary, we have provided evidence that Hipk and caspases engage in a bidirectional positive regulatory relationship that amplifies apoptotic signalling and JNK activation. This molecular crosstalk provides mechanistic insight into a previously reported positive feedback loop between caspase activity and JNK signalling that reinforces the apoptotic fate in *Drosophila* cells (Shlevkov and Morata, 2012). Taken together, prior studies and our current findings delineate a self-reinforcing molecular circuit involving Hipk, JNK signalling, and caspase activation that ensures robust commitment to apoptosis.

## MATERIAL AND METHODS

### Drosophila strains

All the *Drosophila* strains used in this study were raised and maintained on standard medium at 25 °C (see below for the temperature shift experiments). The following *Drosophila* lines were used:

Gal4/UAS and LexA/lexO systems: We have used the Gal4/UAS (Brand and Perrimon, 1993) and lexA/lexO (Lai and Lee, 2006) systems to express or inactivate different genes in particular locations, in some cases combining the two systems so that two adjacent cell populations with distinct genotypes could be compared.

<u>Gal4 lines</u>: *hh-Gal4* (Tanimoto et al., 2000), *tub-Gal80^ts^* (McGuire et al., 2003), *en-Gal4* (BDSC#30564), *Abd-B^LDN^* (*Abd-B*-Gal4^LDN^) (de Navas et al., 2006), *Eip71CD*-Gal4 (Cherbas et al., 2003).

<u>lexO line</u>: *lexO-rpr* (SantaBárbara-Ruiz et al., 2015)

<u>UAS lines</u>: UAS*-miRHG* (Siegrist et al., 2010), UAS*-GFP* (BDSC#5130), UAS-*hid* (Igaki et al., 2000), UAS-*HA-Hipk2M* (Lee et al., 2009), UAS-*HA-Hipk3M* (Lee et al., 2009), UAS-*hipkRNAi* (VDRC KK107857) (Chen and Verheyen, 2012), UAS-*p35* (BDSC#8651), UAS-*cherry* (BDSC#35787), UAS-*lacZ* (BDSC#8529), UAS-*Dronc-GFP-Myc*, MVz-*Drice-Flag-VN*, UAS-*Dronc-GFP-Myc*/MVz-*Drice-Flag-VN* (see below), UAS-*Diap1RNAi* (Leullier et al., 2003), UAS-*droncRNAi* (VDRC #23035).

<u>Mutants</u>: *dronc^i29^*, *dronc^i24^* (Xu et al., 2005).

Reporter lines: *TRE-red* (Chatterjee and Bohmann, 2012).

<u>Construction of the *nub-lexA* transgene</u>. To generate the *nub*-LexA driver line for *nubbin*, we first amplified 3.8 kb of nubbin genomic regulatory DNA (Jenett et al., 2012) using Taq high-fidelity polymerase. The primers used to perform the PCR were:

FP:CACCCTTCAACTTGTAACTGCTGGCTGCA RP:GGGGATTGGTCCGAAAAGAGGATAC PCR products were initially subcloned into the TOPO-TA vector and then transferred as EcoRI fragments into the pBPLexA::GADfluw plasmid (Addgene Plasmid #26232; Pfeiffer et al., 2010). Correct insertion and sequence fidelity were confirmed by Sanger sequencing. Transgenic flies carrying the construct were generated via PhiC31-mediated integration at the attP40 landing site located at cytological position 22F on the second chromosome.

#### Temperature shift experiments

We made use of the Gal4/Gal80^ts^ system (McGuire et al., 2003) to control the time of expression of different genetics constructs. After an egg lay of 1 day at 25°C, larvae including the genetic combinations *hh-Gal4, tub-Gal80^ts^* or *en*-Gal4 *tub*-Gal80^ts^ were raised at 17 °C and then transferred to a restrictive temperature of 29°C or 31°C for 2 or 3 days before dissection. The combined expression of a Gal4 line, *hh*-Gal4 or *en*-Gal4, and *tub*-Gal80^ts^ is represented, for simplicity, as *hh^Gal80^* and *en^Gal80^*, respectively.

### Generation of MVz-Drice-Flag-VN Plasmid

We synthesized a wild-type *Drice* cDNA fused at its C-terminus to a Flag tag and the N-terminal half of a split Venus fluorescent protein, using gene synthesis services provided by Twist Bioscience. The resulting fragment was delivered in a pUC51 plasmid backbone. The *Drice*-Flag-VN construct was then excised from pUC51 as a PmeI–KpnI fragment and subcloned into the corresponding sites of the MVz plasmid (Wendler et al., 2022). Please refer to the plasmid map (Supplementary Fig. 4) for additional details; the full plasmid sequence is available upon request. Transgenic flies carrying the construct were generated via PhiC31-mediated site-specific integration. The construct was inserted at the attP40 site, located at cytological position 25C6.

### Construction of the UAS-Dronc-GFP-Myc

A wild-type *Dronc* cDNA was synthesized (GeneWizz) and fused in-frame to the Suntag and HA-tag peptides at the C-terminal end. To facilitate downstream cloning, additional restriction sites were introduced at both the 5ʹ and 3ʹ ends of the construct, as well as upstream of the tag peptide. The full-length construct was initially subcloned into the pUC57 vector as a *NotI-KpnI* fragment. Subsequently, the vector was digested with *SmaI* and *NheI*, resulting in the removal of the C-terminal Suntag-HA tagging from the wild-type *Dronc* sequence. A modified version of GFP, containing a Myc tag at its C-terminal end, was generated by PCR using the primers listed below. This GFP variant includes a TETDG caspase cleavage site, which, upon *Dronc*-dependent cleavage, restores GFP to a conformational state compatible with fluorescence emission. The template for the GFP-Myc sequence was described previously (Arthurton et al., 2020). The GFP-Myc PCR product was subsequently cloned in-frame at the C-terminal end of wild-type *Dronc* as a *SmaI-NheI* fragment.

Primers used for GFP-Myc amplification:

- Forward primer: 5ʹ GCTTTAATAAGAAACTCTACTTCAATcccgggtttttcaacgaagggggcATGATCAAGATCGCCACCAGGAAGTACC 3ʹ
- Reverse primer: 5ʹ GATAAAATGTCCAGTGGCGGCAAGCTAGCttacaggtcctcctcgctgatcagcttctgctcGTTAGGCAGGTTGTCCACCCTCATCAGG 3ʹ

The complete construct was then subcloned as a *NotI-XhoI* fragment into a UAS-attB *w+* vector previously linearized with *NotI-PspXI*. Please refer to the plasmid map in Supplementary Fig. 4 for further details; full sequence of the plasmid can be distributed upon request.

Transgenic *Drosophila melanogaster* carrying the UAS-Dronc-GFP-TETDG-Myc construct were generated via PhiC31-mediated site-specific integration. The construct was inserted into the attP site located at the 22A3 locus (Bloomington Drosophila Stock Center, stock #9752).

### Construction of the UAS-Dronc-GFP-Myc/MVz-Drice-Flag-VN Dual Plasmid

In parallel with the subcloning of the *Dronc*-GFP-Myc fragment into the standard UAS-attB-white⁺ plasmid, we also inserted this construct into a modified version of UAS-attB-white⁺ in which the loxP site upstream of the UAS repeats had been removed by

NheI digestion followed by re-ligation. The *Dronc*-GFP-Myc fragment was then subcloned as a NotI–XhoI fragment into this modified plasmid, which had been linearized with NotI–PspXI. From the resulting intermediate plasmid, an NsiI–Dronc-GFP-Myc–NsiI fragment was excised and subcloned into the MVz-Drice-Flag-VN plasmid using the same restriction sites. The resulting dual-expression plasmid enables simultaneous expression of *Dronc*-GFP-Myc and *Drice*-Flag-VN under UAS control. A plasmid map is shown in Supplementary Fig. 4, and the full sequence is available upon request. Transgenic flies carrying the construct were generated via PhiC31-mediated integration at the attP40 landing site, located at cytological position 25C6.

### Imaginal discs staining

Third instar larvae were dissected in PBS and fixed with 4% paraformaldehyde, 0.1% deoxycholate (DOC) and 0.3% Triton X-100 in PBS for 27 min at room temperature. They were blocked in PBS, 1% BSA, and 0.3% Triton, incubated with the primary antibody overnight at 4 °C, washed in PBS, 0.3% Triton and incubated with the corresponding fluorescent secondary antibodies for at least 2 h at room temperature in the dark. They were then washed and mounted in Vectashield mounting medium (Vector Laboratories).

The following primary antibodies were used: rat anti-Ci (DSHB 2A1) 1:50; mouse anti-Mmp1 (DSHB, a combination, 1:1:1, of 3B8D12, 3A6B4 and 5H7B11) 1:50; rabbit anti-Hipk (a gift from E. Verheyen) 1:100, rabbit anti-Dcp1 (Cell Signaling, antibody #9578) 1:200, rabbit anti-Diap1 (a gift from H. Steller) 1:2000.

Fluorescently labelled secondary antibodies (Molecular Probes Alexa-488, Alexa-555, Alexa-647, ThermoFisher Scientific) were used in a 1:200 dilution. DAPI (MERCK) and TO-PRO3 (Invitrogen) were used in a 1:1000 dilution to label nuclei. Fluorescently labelled secondary antibodies (Molecular Probes Alexa-488, Alexa-555, Alexa-647, ThermoFisher Scientific) were used in a 1:200 dilution and DAPI (MERCK) was used in a 1:500 dilution to label the nuclei.

### IR treatments

For irradiation experiments, larvae were raised at 17 °C for 3-4 days and then transferred to 31°C 1 day before irradiation. Then, irradiated larvae were grown at 31°C for 3 days before imaginal disc dissection. Larvae were irradiated in an X-ray machine Phillips MG102 at the standard dose of 4000Rads (R).

### Analysis of adult cuticles

Photographs of adult flies were taken with a Leica MZ12 stereomicroscope and a Leica DFC5000 camera, and images were acquired using Leica LAS software (3.7). Theimages were edited and assembled using Photoshop.

### Image acquisition, quantifications and statistical analysis

Stack images were captured with a Leica (Solms, Germany) LSM510, LSM710, DB550 B vertical confocal microscope and a Nikon A1R. Multiple focal planes were obtained for each imaginal disc. Quantifications and image processing were performed using the Fiji/ImageJ (https://fiji.sc) and Adobe Photoshop software.

To measure the percentage of positive area of different markers (%Dcp1, %TREred, %pJNK, %GFP, %Myc), the corresponding positive area was obtained using the “Threshold” tool in ImageJ and then normalized by the area of the compartment (labelled with positive GFP or negative Ci staining). Diap1 ratio (P/A compartment) was calculated as the proportion between the percentage of Diap1 positive areas in the posterior compartment and the anterior compartment.

To quantify the percentage of the posterior compartment, a Z-maximal intensity projection was made for each image. Then, the area of the posterior compartment (labelled with positive GFP or HA staining) was measured by using the “Area” tool and normalized dividing by the total disc area (labelled by TOPRO-3 or DAPI staining).

Flag, Hipk and HA integrated density (ID) were calculated by multiplying the mean intensity (obtained using the “Threshold” tool in ImageJ) and the area of the posterior compartment (labelled by positive Flag, Hipk or HA staining. ID data were normalized by the total disc area (labelled by DAPI or TOPRO staining).

Statistical analysis was performed using the GraphPad Prism v8 software (https://www.graphpad.com). When comparing between two groups, a non-parametric Student’s *t*-test (Mann-Whitney’s test) was used. To compare between more than two groups, a non-parametric, one-way ANOVA test (Kruskal-Wallis test) was used. Sample size was indicated in each graph. Error bars in the graphs represent the standard deviation (SD). p-values obtained in each statistical analysis were represented in the graphs according to the following nomenclature: *p<0,05; **p<0,01; ***p<0,001 and ****p<0,001.

## ACKNOWLEDGMENTS

We thank Raquel Martín for the construction of the *nub*-LexA driver, Esther Verheyen for the anti-Hipk antibody and stocks and Andreas Bergmann for the *dronc* mutants. We also thank the Bloomington Stock Center, the Vienna Drosophila Resource Center, and the Developmental Studies Hybridoma Bank for fly stocks and reagents. This study was supported by grants from FEDER/Ministerio de Ciencia e Innovación-Agencia Estatal de Investigación-Consejo Superior de Investigaciones Científicas [No. PGC2018-095151-B-I00, PID2021-125377NB-100, and PIE Intramural 202020E255 to GM, BFU2017-86244-P and PID2020-113318GB-I00 to ES, and PID2023-150773NB-100 to LAB). J.M. G. was a recipient of a Formación del Personal Investigador (FPI) fellowship (PRE 2019_090108) from the Spanish Government and R.A. J. was a recipient of a CONACyT Fellowship. Institutional support from the Ramón Areces Foundation is acknowledged.

## Supplementary Figures

**Supplementary Figure 1.**
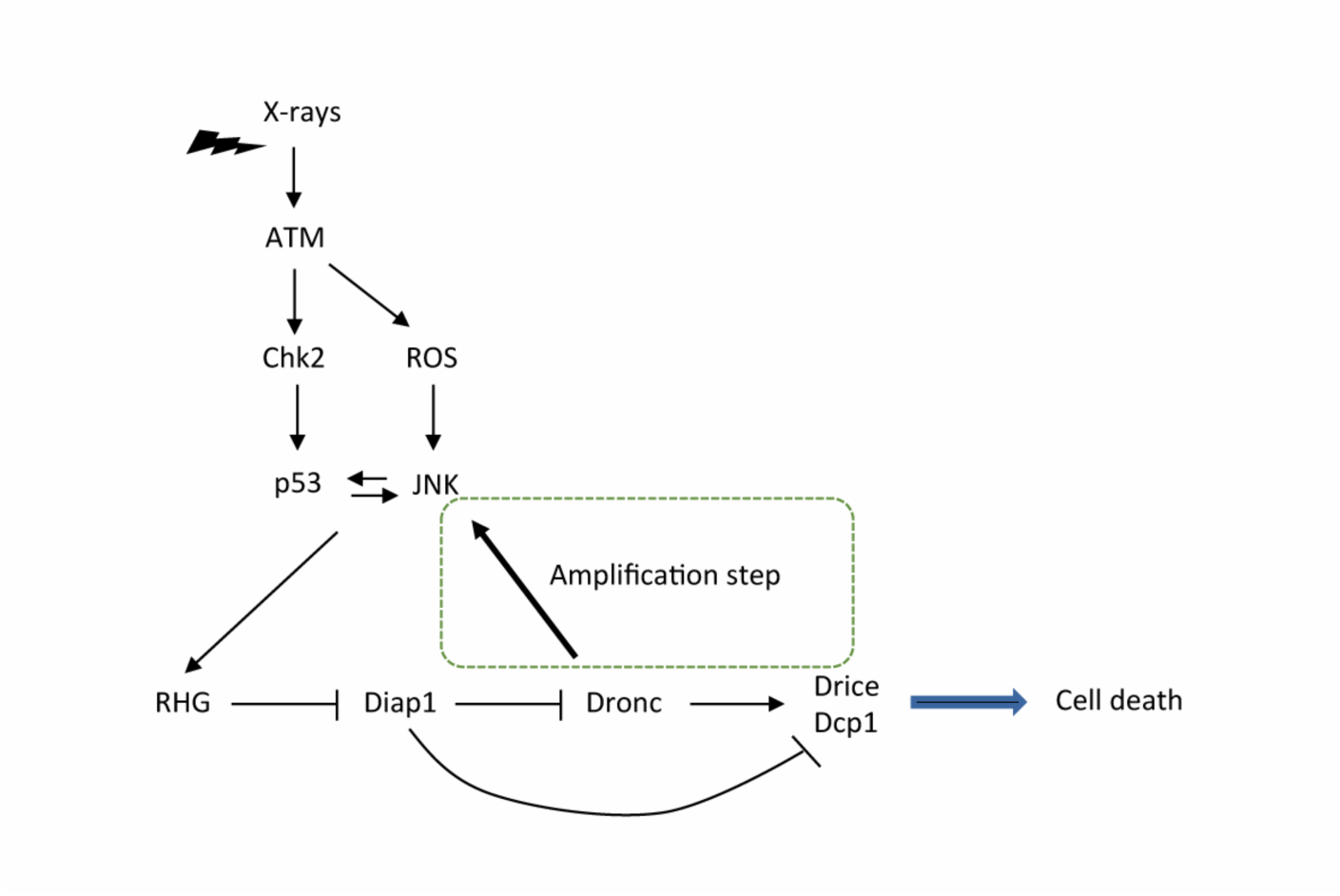
**Simplified version of the apoptosis program of *Drosophila*, as triggered by X-rays.** The DNA double-strand breaks caused by the irradiation activates the Ataxia Telengiestasia Mutated (ATM) kinase, which in turn activates the Checkpoint2 (Chk2) kinase and also induces the production of Reactive Oxygen Species (ROS). These events trigger the function of *p53* and of the Jun N-Terminal Kinase (JNK) pathway, known to stimulate each other. The expression of *p53*/JNK transcriptionally activates the pro-apoptotic genes *reaper (rpr), head involution defective (hid)* and *grim*, which cause ubiquitination of the Drosophila inhibitor of apoptosis protein1 (Diap1) and allow activation of the apical caspase Dronc and subsequently of the effector caspases Drice and Dcp1. In addition of activating the effector caspases, Dronc stimulates JNK levels, thus establishing an amplification loop, necessary for the full apoptotic response. The amplification step is squared for emphasis.

**Supplementary Figure 2.**
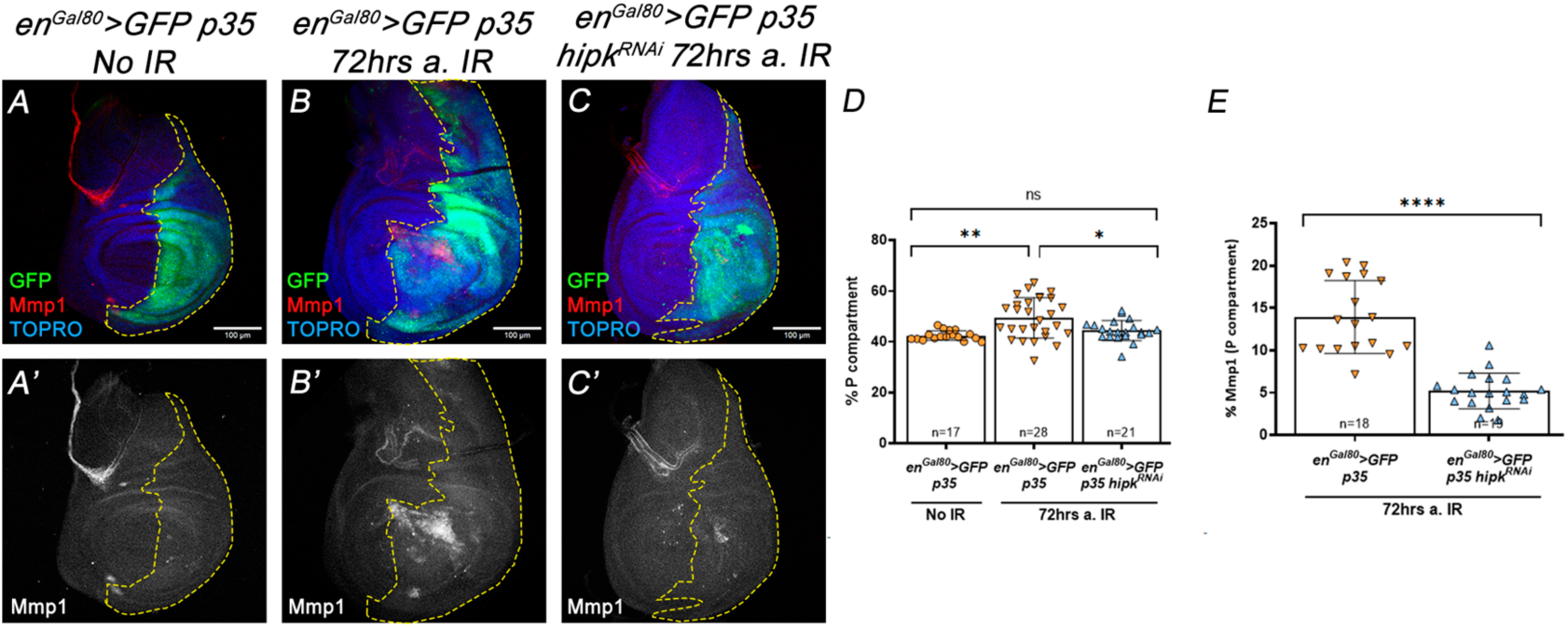
***hipk* is required to maintain JNK function and size increase after irradiation (IR)** Genotypes on top of the panels. (A, A’) Non-irradiated discs show no JNK activity, monitored here by the presence of Metalloprotease 1, Mmp1 (in red), a target of JNK. (B, B’) In irradiated discs of the *en^Gal80^*>*GFP p35* genotype, the presence of the baculovirus protein P35 in the P compartment allows the survival of cells in which JNK has been induced by IR, thus increasing Mmp1 signal. Size of the compartment is also increased. (C, C’) In *p35*-expressing irradiated discs in which *hipk* function is reduced by the expression of a *hipk^RNAi^* construct, JNK activity and compartment size are much diminished. Quantifications in D, E.

**Supplementary Figure 3.**
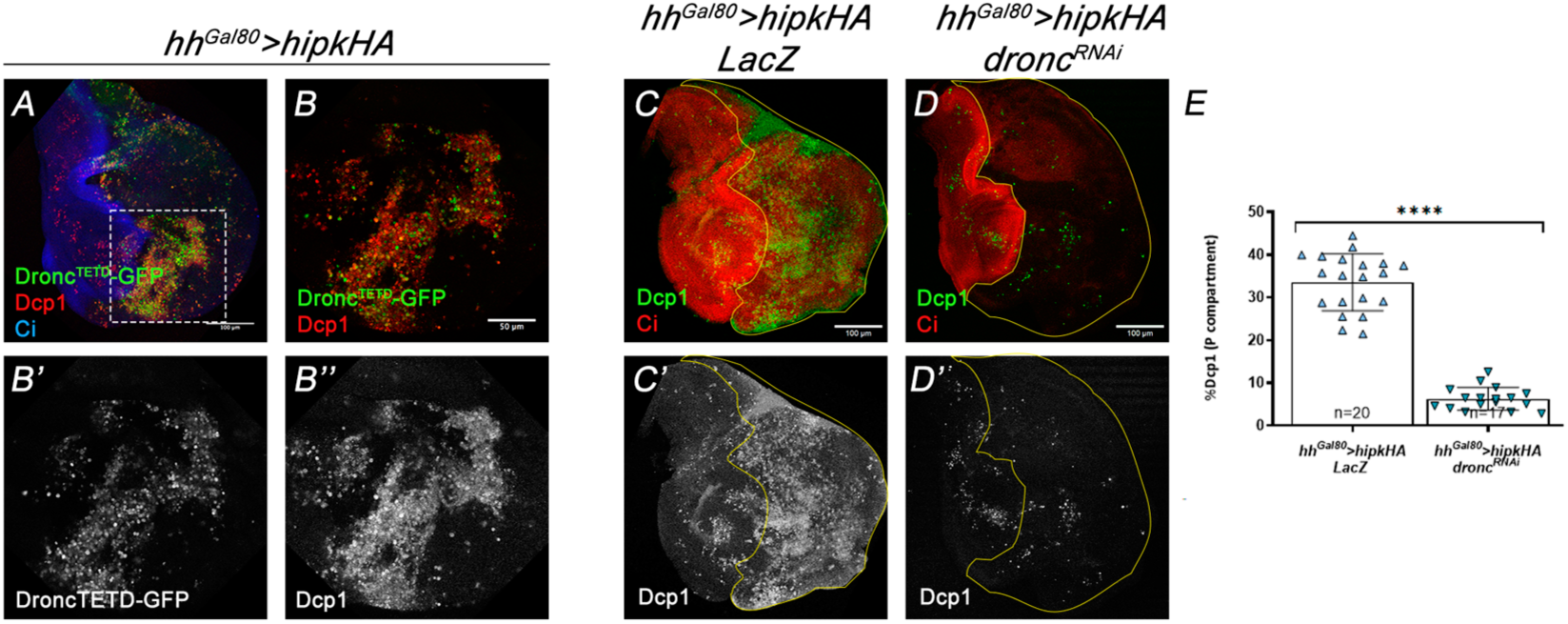
**Overexpression of *hipk* results in *dronc*-dependent apoptosis** Genotypes on top of the panel. (A-B’’) Overexpression of *hipk-HA* induces the expression of the Dronc-activity reporter *dronc^TETD^-GFP* and Dcp1 antibody signal (A). B-B’’ are amplifications of the inset in A. (C-D’’) The Dcp1 signal induced by overexpression of *hipk-HA* (C, C’) is significantly reduced by the simultaneous inactivation of *dronc* (D, D’). Ci, in red, marks the A compartment (in blue in A, in red in C, D). Quantifications in E.

**Supplementary Figure 4.**
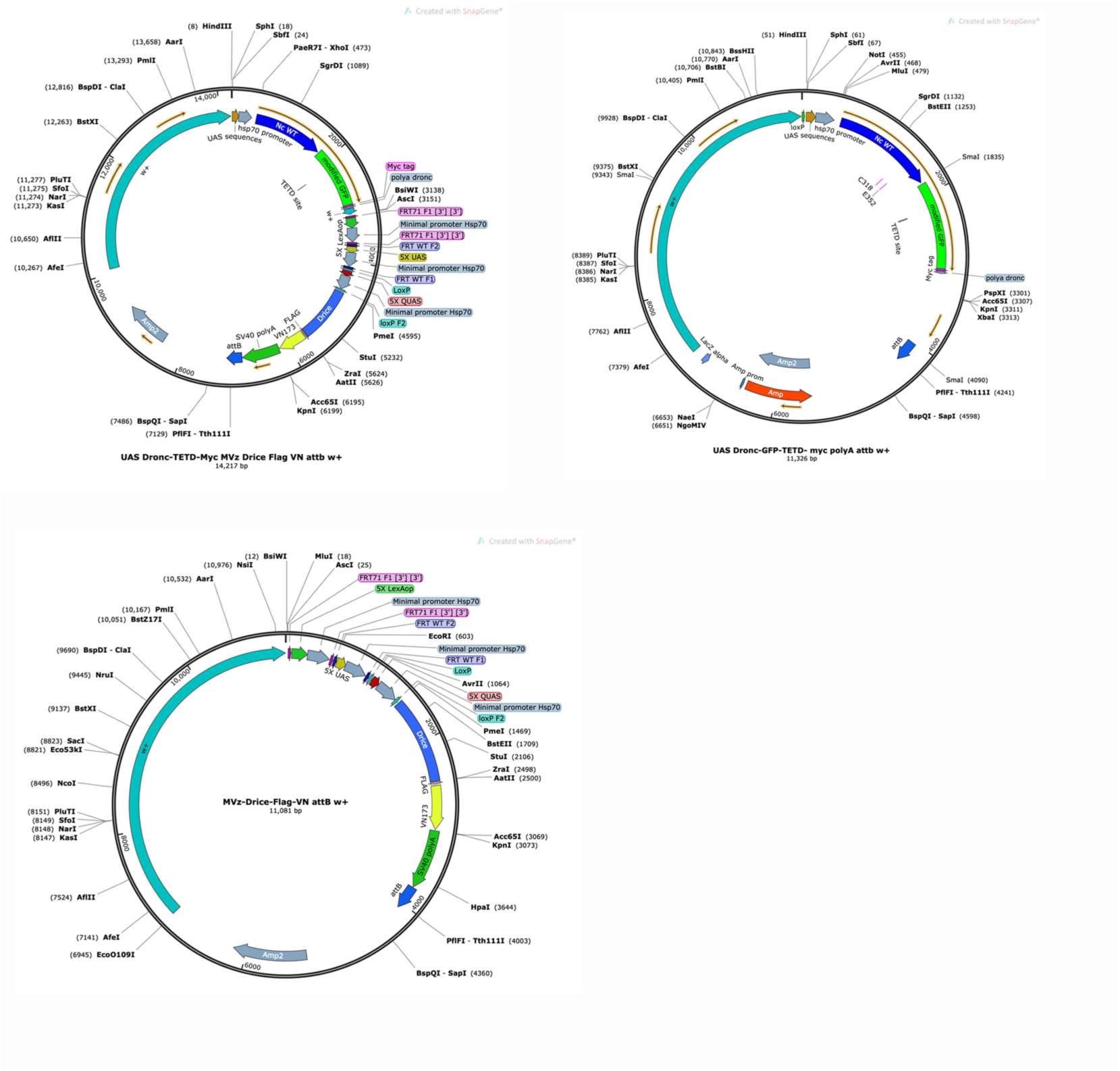
**Plasmid maps of the MVz-Drice-Flag-VN, UAS-Dronc-GFP-Myc and UAS-Dronc-GFP-Myc/MVz-Drice-Flag-VN plasmids.**

## REFERENCES

Abbott MK, Lengyel JA. Embryonic head involution and rotation of male terminalia require the Drosophila locus head involution defective. Genetics. 1991 Nov;129(3):783–9. doi: 10.1093/genetics/129.3.783.

Allan LA, Clarke PR. Apoptosis and autophagy: Regulation of caspase-9 by phosphorylation. FEBS J. 2009 Nov;276(21):6063–73. doi: 10.1111/j.1742-4658.2009.07330.x.

Amcheslavsky A, Wang S, Fogarty CE, Lindblad JL, Fan Y, Bergmann A. Plasma Membrane Localization of Apoptotic Caspases for Non-apoptotic Functions Dev Cell. 2018 May 21;45(4):450–464.e3. doi: 10.1016/j.devcel.2018.04.020.

Arthurton L, Nahotko DA, Alonso J, Wendler F, Baena-Lopez LA. Non-apoptotic caspase activation preserves Drosophila intestinal progenitor cells in quiescence EMBO Rep. 2020 Dec 3;21(12):e48892. doi: 10.15252/embr.201948892.

Baonza, A., Tur-Gracia, S., Pérez-Aguilera, M., Estella, C. Regulation and coordination of the different DNA damage responses in Drosophila Front. Cell Dev. Biol. 10:993257. doi: 10.3389/fcell.2022.993257.

Bischoff M, Cseresnyés Z. Cell rearrangements, cell divisions and cell death in a migrating epithelial sheet in the abdomen of Drosophila. Development 2009. 136, 2403–11. doi: 10.1242/dev.035410.

Blaquiere JA, Verheyen EM. Homeodomain-Interacting Protein Kinases: Diverse and Complex Roles in Development and Disease. Curr Top Dev Biol. 2017;123:73–103. doi: 10.1016/bs.ctdb.2016.10.002.

Blaquiere JA, Wong KKL, Kinsey SD, Wu J, Verheyen EM. Homeodomain-interacting protein kinase promotes tumorigenesis and metastatic cell behavior. Dis Model Mech. 2018 Jan 17;11(1): dmm031146. doi: 10.1242/dmm.031146.

Brady SC, Allan LA, Clarke PR. Regulation of caspase 9 through phosphorylation by protein kinase C zeta in response to hyperosmotic stress. Mol Cell Biol. 2005 Dec;25(23):10543–55. doi: 10.1128/MCB.25.23.10543-10555.2005.

Brand AH, Perrimon N. Targeted gene expression as a means of altering cell fates and generating dominant phenotypes. Development. 1993 Jun;118(2):401–15. doi: 10.1242/dev.118.2.401.

Chatterjee N, Bohmann D. A versatile ΦC31 based reporter system for measuring AP-1 and Nrf2 signaling in Drosophila and in tissue culture. PLoS One. 2012;7(4):e34063. doi: 10.1371/journal.pone.0034063.

Chen P, Nordstrom W, Gish B, Abrams JM. grim, a novel cell death gene in Drosophila. Genes Dev. 1996 Jul 15;10(14):1773–82. doi: 10.1101/gad.10.14.1773.

Chen J, Verheyen EM. Homeodomain-interacting protein kinase regulates Yorkie activity to promote tissue growth. Curr Biol. 2012 Sep 11;22(17):1582–6. doi: 10.1016/j.cub.2012.06.074.

Cherbas L, Hu X, Zhimulev I, Belyaeva E, Cherbas P. EcR isoforms in Drosophila: testing tissue-specific requirements by targeted blockade and rescue. Development. 2003 Jan;130(2):271–84. doi: 10.1242/dev.00205.

Choi CY, Kim YH, Kim YO, Park SJ, Kim EA, Riemenschneider W, Gajewski K, Schulz RA, Kim Y. Phosphorylation by the DHIPK2 protein kinase modulates the corepressor activity of Groucho. J Biol Chem. 2005 Jun 3;280(22):21427–36. doi: 10.1074/jbc.M500496200.

Dauth I, Krüger J, Hofmann TG. Homeodomain-interacting protein kinase 2 is the ionizing radiation-activated p53 serine 46 kinase and is regulated by ATM. Cancer Res. 2007 Mar 1;67(5):2274–9. doi: 10.1158/0008-5472.CAN-06-

D’Brot A, Chen P, Vaishnav M, Yuan S, Akey CW, Abrams JM. Tango7 directs cellular remodeling by the Drosophila apoptosome. Genes Dev. 2013 Aug 1;27(15):1650–5. doi: 10.1101/gad.219287.113.

de Navas L, Foronda D, Suzanne M, Sánchez-Herrero E. A simple and efficient method to identify replacements of P-lacZ by P-Gal4 lines allows obtaining Gal4 insertions in the bithorax complex of Drosophila. Mech Dev. 2006 Nov;123(11):860–7. doi: 10.1016/j.mod.2006.07.010.

Di Stefano V, Rinaldo C, Sacchi A, Soddu S, D’Orazi G. Homeodomain-interacting protein kinase-2 activity and p53 phosphorylation are critical events for cisplatin-mediated apoptosis. Exp Cell Res. 2004 Feb 15;293(2):311–20. doi: 10.1016/j.yexcr.2003.09.032.

Dorstyn L, Colussi PA, Quinn LM, Richardson H, Kumar S. Proc Natl Acad Sci U S A. DRONC, an ecdysone-inducible Drosophila caspase. 1999 Apr 13;96(8):4307–12. doi: 10.1073/pnas.96.8.4307.

Fogarty CE, Bergmann A. Detecting caspase activity in Drosophila larval imaginal discs. Methods Mol Biol. 2014;1133:109–17. doi: 10.1007/978-1-4939-0357-3_7.

Fogarty CE, Diwanji N, Lindblad JL, Tare M, Amcheslavsky A, Makhijani K, Brückner K, Fan Y, Bergmann A. Extracellular Reactive Oxygen Species drive Apoptosis-induced Proliferation via *Drosophila* Macrophages. Curr Biol. 2016 Mar 7;26(5):575–84. doi: 10.1016/j.cub.2015.12.064.

Fraser AG, Evan GI. Identification of a Drosophila melanogaster ICE/CED-3-related protease, drICE. EMBO J. 1997 May 15;16(10):2805–13. doi: 10.1093/emboj/16.10.2805.

Fuchs Y, Steller H. Programmed cell death in animal development and disease. Cell. 2011 Nov 11;147(4):742–58. doi: 10.1016/j.cell.2011.10.033.

Ge J, Wang Y, Li X, Lu Q, Yu H, Liu H, Ma K, Deng X, Luo ZQ, Liu X, Qiu J. Phosphorylation of caspases by a bacterial kinase inhibits host programmed cell death Nat Commun. 2024 Sep 30;15(1):8464. doi: 10.1038/s41467-024-52817-1.

Gleichauf, R. Anatomie und variabilität des geslechtsapparates von D. Melanogaster Meigen. Z. wiss Zool., 148 (1936), pp. 1-66.

Goyal L, McCall K, Agapite J, Hartwieg E, Steller H . Induction of apoptosis by *Drosophila* reaper, hid and grim through inhibition of IAP function. EMBO J 2000; 19: 589–597. doi: 10.1093/emboj/19.4.589.

Grether ME, Abrams JM, Agapite J, White K, Steller H. The head involution defective gene of Drosophila melanogaster functions in programmed cell death. Genes Dev. 1995 Jul 15;9(14):1694–708. doi: 10.1101/gad.9.14.1694.

Gresko E, Roscic A, Ritterhoff S, Vichalkovski A, del Sal G, Schmitz ML. Autoregulatory control of the p53 response by caspase-mediated processing of HIPK2. EMBO J. 2006 May 3;25(9):1883–94. doi: 10.1038/sj.emboj.7601077.

Hay BA, Wolff T, Rubin GM. Expression of baculovirus P35 prevents cell death in Drosophila. Development. 1994 Aug;120(8):2121–9. doi: 10.1242/dev.120.8.2121.

Hay BA, Wassarman DA, Rubin GM. Drosophila homologs of baculovirus inhibitor of apoptosis proteins function to block cell death. Cell. 1995 Dec 29;83(7):1253–62. doi: 10.1016/0092-8674(95)90150-7.

Hofmann TG, Stollberg N, Schmitz ML, Will H. HIPK2 regulates transforming growth factor-beta-induced c-Jun NH(2)-terminal kinase activation and apoptosis in human hepatoma cells. Cancer Res. 2003 Dec 1;63(23):8271–7.

Huang H, Du G, Chen H, Liang X, Li C, Zhu N, Xue L, Ma J, Jiao R. Drosophila Smt3 negatively regulates JNK signaling through sequestering Hipk in the nucleus. Development. 2011 Jun;138(12):2477–85. doi: 10.1242/dev.061770.

Igaki, T., Kanuka, H., Inohara, N., Sawamoto, K., Nunez, G., Okano, H., & Miura, M. Drob-1, a Drosophila member of the Bcl-2/CED-9 family that promotes cell death. Proceedings of the National Academy of Sciences. 2000; 97(2),662–667. 10.1073/pnas.97.2.662

Igaki, T., Kanda, H., Yamamoto-Goto, Y., Kanuka, H., Kuranaga, E., Aigaki, T., & Miura, M. Eiger, a TNF superfamily ligand that triggers the Drosophila JNK pathway. The EMBO Journal. 2002; 21(12), 3009–3018. doi: 10.1093/emboj/cdf306.

Jacobsen MD, Weil M, Raff MC. Role of Ced-3/ICE-family proteases in staurosporine-induced programmed cell death. J Cell Biol. 1996;133(5):1041–51).

Jenett A, Rubin GM, Ngo TT, Shepherd D, Murphy C, Dionne H, Pfeiffer BD, Cavallaro A, Hall D, Jeter J, Iyer N, Fetter D, Hausenfluck JH, Peng H, Trautman ET, Svirskas RR, Myers EW, Iwinski ZR, Aso Y, DePasquale GM, Enos A, Hulamm P, Lam SC, Li HH, Laverty TR, Long F, Qu L, Murphy SD, Rokicki K, Safford T, Shaw K, Simpson JH, Sowell A, Tae S, Yu Y, Zugates CT. A GAL4-driver line resource for Drosophila neurobiology. Cell Rep. 2012 Oct 25;2(4):991–1001. doi: 10.1016/j.celrep.2012.09.011.

Kester RS, Nambu JR. Targeted expression of p35 reveals a role for caspases in formation of the adult abdominal cuticle in Drosophila. Int J Dev Biol. 2011;55(1):109–19. doi: 10.1387/ijdb.103109rk.

Kuranaga E., Matsunuma T., Kanuka H., Takemoto K., Koto A., Kimura K., Miura M. Apoptosis controls the speed of looping morphogenesis in *Drosophila* male terminalia. Development 2011; 138, 1493–99.

Lai SL, Lee T. Genetic mosaic with dual binary transcriptional systems in Drosophila. Nat Neurosci. 2006;9:703–709. doi: 10.1038/nn1681

Lee, W., Swarup, S., Chen, J., Ishitani, T., & Verheyen, E. M. Homeodomain interacting protein kinases (Hipks) promote Wnt/Wg signaling through stabilization of -catenin/Arm and stimulation of target gene expression. Development.2009a. 136(2), 241–251. doi: 10.1016/j.ydbio.2008.10.029.

Lee W, Andrews BC, Faust M, Walldorf U, Verheyen EM. Hipk is an essential protein that promotes Notch signal transduction in the Drosophila eye by inhibition of the global co-repressor Groucho. Dev Biol. 2009b Jan 1;325(1):263–72. doi: 10.1016/j.ydbio.2008.10.029.

Leulier F, Ribeiro PS, Palmer E, Tenev T, Takahashi K, Robertson D, Zachariou A, Pichaud F, Ueda R, Meier P. Systematic in vivo RNAi analysis of putative components of the Drosophila cell death machinery. Cell Death Differ. 2006 Oct;13(10):1663–74. doi: 10.1038/sj.cdd.4401868.

Link N, Chen P, Lu WJ, Pogue K, Chuong A, Mata M, Checketts J, Abrams JM. A collective form of cell death requires homeodomain interacting protein kinase J Cell Biol. 2007 Aug 13;178(4):567–74. doi: 10.1083/jcb.200702125.

Lisi S, Mazzon I, White K. Diverse domains of THREAD/DIAP1 are required to inhibit apoptosis induced by REAPER and HID in Drosophila. Genetics. 2000 Feb;154(2):669–78. doi: 10.1093/genetics/154.2.669.

Liu J, Lin A. Role of JNK activation in apoptosis: a double-edged sword. Cell Res. 2005;15:36–42. doi: 10.1038/sj.cr.7290262.

Lohmann I, McGinnis N, Bodmer M, McGinnis W. “The Drosophila Hox gene deformed sculpts head morphology via direct regulation of the apoptosis activator reaper Cell. 110, 457–66 (2002) doi: 10.1016/s0092-8674(02)00871-1.

Macías A, Romero NM, Martín F, Suárez L, Rosa AL, Morata G. PVF1/PVR signaling and apoptosis promotes the rotation and dorsal closure of the Drosophila male terminalia. Int J Dev Biol. 2004 Dec;48(10):1087–94. doi: 10.1387/ijdb.041859am.

Madhavan MM, Madhavan K. Morphogenesis of the epidermis of adult abdomen of Drosophila. J Embryol Exp Morphol. 1980 Dec;60:1–31.

Martin MC, Allan LA, Lickrish M, Sampson C, Morrice N, Clarke PR. Protein kinase A regulates caspase-9 activation by Apaf-1 downstream of cytochrome c. J Biol Chem. 2005 Apr 15;280(15):15449–55. doi: 10.1074/jbc.M414325200.

Martín R, Pinal N, Morata G. Distinct regenerative potential of trunk and appendages of *Drosophila* mediated by JNK signalling. Development. 2017 Nov 1;144(21):3946–3956. doi: 10.1242/dev.155507.

McGuire SE, Le PT, Osborn AJ, Matsumoto K, Davis RL. Spatiotemporal rescue of memory dysfunction in Drosophila. Science. 2003;302:1765–8. doi: 10.1126/science.1089035.

McNamee, L., Brodsky, M. p53-Independent Apoptosis Limits DNA Damage-Induced Aneuploidy. Genetics. 2009 Jun;182(2):423–35. doi: 10.1534/genetics.109.102327.

McEwen DG, Peifer M. Puckered, a Drosophila MAPK phosphatase, ensures cell viability by antagonizing JNK-induced apoptosis. 2005 Sep;132(17):3935–46. doi: 10.1242/dev.01949. Development 132: 3935–3946.

Meier P, Silke J, Leevers SJ, Evan GI. The Drosophila caspase DRONC is regulated by DIAP1. EMBO J. 2000 Feb 15;19(4):598–611. doi: 10.1093/emboj/19.4.598.

Nakajima Y, Kuranaga E, Sugimura K, Miyawaki A, Miura M. Nonautonomous apoptosis is triggered by local cell cycle progression during epithelial replacement in Drosophila. Mol Cell Biol. 2011 Jun;31(12):2499–512. doi: 10.1128/MCB.01046-10.

Ninov N, Chiarelli DA, Martín-Blanco E. Extrinsic and intrinsic mechanisms directing epithelial cell sheet replacement during Drosophila metamorphosis. Development. 2007 Jan;134(2):367–79. doi: 10.1242/dev.02728

Pérez-Garijo A, Martín FA, Morata G. Caspase inhibition during apoptosis causes abnormal signalling and developmental aberrations in Drosophila. Development. 2004 Nov;131(22):5591–8. doi: 10.1242/dev.01432.

Pfeiffer, et al., 2010. Pfeiffer BD, Ngo TT, Hibbard KL, Murphy C, Jenett A, Truman JW, Rubin GM. Refinement of tools for targeted gene expression in Drosophila. Genetics. 2010 Oct;186(2):735-55. doi: 10.1534/genetics.110.119917.

Pinal N, Martín M, Medina I, Morata G. Short-term activation of the Jun N-terminal kinase pathway in apoptosis-deficient cells of Drosophila induces tumorigenesis Nat Commun. 2018 Apr 18;9(1):1541. doi: 10.1038/s41467-018-04000-6.

Pinal N, Calleja M, Morata G. Pro-apoptotic and pro-proliferation functions of the JNK pathway of Drosophila: roles in cell competition, tumorigenesis and regeneration Open Biol. 2019 Mar 29;9(3):180256. doi: 10.1098/rsob.180256.

Poon CL, Zhang X, Lin JI, Manning SA, Harvey KF. Homeodomain-interacting protein kinase regulates Hippo pathway-dependent tissue growth. Curr Biol. 2012 Sep 11;22(17):1587–94. doi: 10.1016/j.cub.2012.06.075.

Rinaldo C, Prodosmo A, Siepi F, Soddu S. HIPK2: a multitalented partner for transcription factors in DNA damage response and development. Biochem Cell Biol. 2007 Aug;85(4):411–8. doi: 10.1139/O07-071.

Rodriguez A, Oliver H, Zou H, Chen P, Wang X, Abrams JM. Dark is a Drosophila homologue of Apaf-1/CED-4 and functions in an evolutionarily conserved death pathway Nat Cell Biol. 1999 Sep;1(5):272–9. doi: 10.1038/12984.

Ryoo HD, Bergmann A, Gonen H, Ciechanover A, Steller H. Regulation of Drosophila IAP1 degradation and apoptosis by reaper and ubcD1. Nat Cell Biol. 2002 Jun; 4(6):432–8. doi: 10.1038/ncb795.

Ryoo HD, Gorenc T, Steller. Apoptotic cells can induce compensatory cell proliferation through the JNK and the Wingless signaling pathways. Dev Cell. 2004 Oct;7(4):491–501. doi: 10.1016/j.devcel.2004.08.019.

Santabárbara-Ruiz P, López-Santillán M, Martínez-Rodríguez I, Binagui-Casas A, Pérez L, Milán M, Corominas M, Serras F. ROS-Induced JNK and p38 Signaling Is Required for Unpaired Cytokine Activation during Drosophila Regeneration. PLoS Genet. 2015 Oct 23;11(10):e1005595. doi: 10.1371/journal.pgen.1005595.

Schmitz ML, Rodriguez-Gil A, Hornung Integration of stress signals by homeodomain interacting protein kinases. J. Biol Chem. 2014 Apr;395(4):375–86. doi: 10.1515/hsz-2013-0264.

Shang Y, Zhang J, Huang EJ. HIPK2-Mediated Transcriptional Control of NMDA Receptor Subunit Expression Regulates Neuronal Survival and Cell Death. J Neurosci. 2018 Apr 18;38(16):4006–4019. doi: 10.1523/JNEUROSCI.3577-17.2018.

Shlevkov E, Morata G. A dp53/JNK-dependant feedback amplification loop is essential for the apoptotic response to stress in Drosophila. Cell Death Differ. 2012 Mar;19(3):451–60. doi: 10.1038/cdd.2011.113.

Siegrist, S. E., N. S. Haque, C. H. Chen, B. A. Hay and I. K. Hariharan (2010). Inactivation of both Foxo and 975 reaper promotes long-term adult neurogenesis in Drosophila. Curr Biol Apr 13;20(7):643–8. doi: 10.1016/j.cub.2010.01.060.

Smith-Bolton RK, Worley MI, Kanda H, Hariharan IK. Regenerative growth in Drosophila imaginal discs is regulated by Wingless and Myc. Dev Cell. 2009 Jun;16(6):797–809. doi: 10.1016/j.devcel.2009.04.015.

Sombroek D, Hofmann TG. How cells switch HIPK2 on and off. Cell Death Differ. 2009 Feb;16(2):187–94. doi: 10.1038/cdd.2008.154

Spéder P, Adám G, Noselli S. Type ID unconventional myosin controls left-right asymmetry in Drosophila. Nature. 2006 Apr 6;440(7085):803-7. doi: 10.1038/nature04623.

Steinmetz EL, Dewald DN, Walldorf U. *Drosophila* Homeodomain-Interacting Protein Kinase (Hipk) Phosphorylates the Hippo/Warts Signalling Effector Yorkie. Int J Mol Sci. 2021 Feb 13;22(4):1862. doi: 10.3390/ijms22041862.

Steller H. Regulation of apoptosis in Drosophila. Cell Death Differ 2008 Jul;15(7):1132–8. doi: 10.1038/cdd.2008.50.

Suzanne M., Pedtzoldt A.G., Spéder P., Coutelis J.B., Steller H., Noselli S. Coupling of apoptosis and L/R patterning controls stepwise organ looping. Current Biology 2010; 20, 1773–78.

Tanimoto H, Itoh S, ten Dijke P, Tabata T. Hedgehog creates a gradient of DPP activity in Drosophila wing imaginal discs. Mol cell. 2000;5:59–71. doi: 10.1016/S1097-2765(00)80403-7.

Tettweiler G, Blaquiere JA, Wray NB, Verheyen EM. Hipk is required for JAK/STAT activity during development and tumorigenesis. PLoS One. 2019 Dec 31;14(12):e0226856. doi: 10.1371/journal.pone.0226856.

Uhlirova M, Bohmann D. JNK-and Fos-regulated Mmp1 expression cooperates with Ras to induce invasive tumors in Drosophila. EMBO J. 2006 Nov 15;25(22):5294–304. doi: 10.1038/sj.emboj.7601401.

Verghese S, Su TT. Drosophila Wnt and STAT Define Apoptosis-Resistant Epithelial Cells for Tissue Regeneration after Irradiation. PLoS Biol. 2016 Sep 1;14(9):e1002536. doi: 10.1371/journal.pbio.1002536.

Verheyen EM, Swarup S, Lee W. Hipk proteins dually regulate Wnt/Wingless signal transduction. Fly (Austin). 2012 Apr-Jun;6(2):126-31. doi: 10.4161/fly.20143.

Wang SL, Hawkins CJ, Yoo SJ, Müller HA, Hay BA. The Drosophila caspase inhibitor DIAP1 is essential for cell survival and is negatively regulated by HID. Cell. 1999 Aug 20;98(4):453–63. doi: 10.1016/s0092-8674(00)81974-1.

Wang Y, Debatin KM, Hug H. HIPK2 overexpression leads to stabilization of p53 protein and increased p53 transcriptional activity by decreasing Mdm2 protein levels. BMC Mol Biol. 2001;2:8. doi: 10.1186/1471-2199-2-8.

Wells BS, Yoshida E, Johnston LA. Compensatory proliferation in Drosophila imaginal discs requires Dronc-dependent p53 activity. Curr Biol. 2006 Aug 22;16(16):1606–15. doi: 10.1016/j.cub.2006.07.046.

Wendler F, Park S, Hill C, Galasso A, Chang KR, Awan I, Sudarikova Y, Bustamante-Sequeiros M, Liu S, Sung EY, Aisa-Bonoko G, Kim SK, Baena-Lopez LA. A LexAop > UAS > QUAS trimeric plasmid to generate inducible and interconvertible Drosophila overexpression transgenes. Sci Rep. 2022 Mar 9;12(1):3835. doi: 10.1038/s41598-022-07852-7.

Weston CR, Davis RJ. The JNK signal transduction pathway. Curr Opin Cell Biol. 2007 Apr;19(2):142–9. doi: 10.1016/j.ceb.2007.02.001.

White K, Tahaoglu E, Steller H. Science. Cell killing by the Drosophila gene reaper. 1996 Feb 9;271(5250):805-7. doi: 10.1126/science.271.5250.805.

Yang CS, Thomenius MJ, Gan EC, Tang W, Freel CD, Merritt TJ, Nutt LK, Kornbluth S. Metabolic regulation of Drosophila apoptosis through inhibitory phosphorylation of Dronc EMBO J. 2010 Sep 15;29(18):3196–207. doi: 10.1038/emboj.2010.191.

Xu D, Li Y, Arcaro M, Lackey M, Bergmann A. The CARD-carrying caspase Dronc is essential for most, but not all, developmental cell death in Drosophila. Development. 2005 May;132(9):2125–34. doi: 10.1242/dev.01790.

Zhang Q, Yoshimatsu Y, Hildebrand J, Frisch SM, Goodman RH. Homeodomain interacting protein kinase 2 promotes apoptosis by downregulating the transcriptional corepressor CtBP. <cell. 2003 Oct 17;115(2):177–86. doi: 10.1016/s0092-8674(03)00802-x.

